# *Medicago Sativa* Defensin 1 (MsDef1), A Natural Tumor Targeted Sensitizer for Improving Chemotherapy: Translation from Anti-Fungal Agent to Potential Anti-Cancer Agent

**DOI:** 10.1101/2021.02.13.431112

**Authors:** Raghu S. Pandurangi, Amol Karwa, Uma Shankar Sagaram, Dilip Shah

## Abstract

**Introduction:** MsDef1, a 45-amino acid cysteine-rich peptide from the seed of *Medicago sativa* is an antifungal defensin small protein. It exhibits broad-spectrum antifungal activity against fungal pathogens of plants at low micromolar concentrations. The common vulnerability of fungal and cancer cells determines the utility of MsDef1 as a potential anti-tumor agent.

**Results:** The solution dynamics of ^15^N-labeled MsDef1, ^15^N longitudinal relaxation (T1) and ^15^N-^1^H Nuclear Overhauser Effect (NOE) shows that GlcCer binds at two sites on the peptide molecule, i.e., Asp36-Cys39 and amino acids between 12-20 and 33-40. MsDef1 interacts with drug resistant breast cancer MCF-7R cells, permeates GlcCer-rich plasma membrane and releases apoptotic ceramide. This results in the activation of ceramide pathway involving interaction of the peptide with intracellular thioredoxin (Trx), another tumor specific biomarker. MsDef1 oxidizes Trx through four S-S bonds and in the process, gets reduced to thiols. Oxidation of Trx is correlated with the activation of Apoptosis Stimulating Kinase 1 (ASK1) which is known to sensitize cancer cells to chemotherapeutics including front-line drug Doxorubicin. A combination of MsDef1 and Doxorubicin exhibits 5-10-fold greater apoptosis *in vitro* in MDR triple negative breast cancer (TNBC) cells compared to either MsDef1 or Doxorubicin alone.

**Conclusion:** An antifungal plant defensin MsDef1 is shown to be a cell permeating peptide (CPP) for MDR cancer cells targeted to two tumor specific targets activating two cell death pathways. That makes MsDef1, potentially a tumor targeted sensitizer neoadjuvant to cancer therapy.

## Introduction

The development of multidrug resistance (MDR) is a serious clinical problem responsible for therapy failure, relapse of cancer and making tumors refractory to future treatments^1^. Deactivation of cell death pathways (e.g., ceramide, apoptosis stimulating kinase (ASK1)^2^ and avoidance of immune surveillance^3^ play major roles in desensitizing cancer cells to treatments of varying nature ^3b^. As a result, high drug dose is needed to treat the cancer which in turn induces stemness into cancer cells, increases resistance, suppresses immune function and enhances off-target toxicity leading to side effects^4^.

Despite its moderate to low efficacy and high off-target toxicity, chemotherapy is still the front-line treatment for most cancers^5^. The major problem with chemotherapy is that it kills only bulk non-stem cancer cells leaving behind resistant cells. This results in high disease recurrence (13% for kidney cancer, 36% for breast and almost 100% for brain cancer^6^) and makes tumors refractory to future treatments^7^. Dose related side effects such as acute lymphedema and low platelets may force discontinuation of treatment altogether creating more resistant cancer cells^8^ thus, causing vicious circles. Sensitizing resistant tumor cells is known to evoke a better response from chemotherapy^9^. However, sensitizing tumor cells <u>selectively</u> using multi drug resistant (MDR) biomarkers is of prime importance in order to make chemotherapy effective in lowering the side effects. This is true particularly for triple negative breast cancer (TNBC) patients who have no option but chemotherapy with a >90% recurrence rate and a median survival of 13 months^10^. The situation is worse for African Americans with recurrence rates of >95%. Current targeted treatments for TNBC patients do not work since they lack biomarkers (e.g., ER, PR and HER2 negative) for which drugs are designed^11^. That leaves non-specific anthracyclines (e.g., Doxorubicin, paclitaxel) as sole options in spite of their high side effects and suppressed immune function^12^.

Cancer cells desensitize themselves to chemotherapy for their survival. For example, TNBC cells circumvent Doxorubicin effect by glycosylating lethal, apoptotic ceramide using glucosylceramide synthase enzyme (GCS) to non-apoptotic glucosylceramide (GlcCer) which becomes a <u>biomarker</u> of MDR (Scheme 1)^13^. Similarly, GlcCer is consistently present at high levels in <u>drug resistant tumors</u> and in tumors taken from patients who are non-responsive to chemotherapy^14^. GlcCer was low in patients who responded to chemotherapy^15^. Stemness of cancer cells increases as glycosylation increases^16^. In fact, ceramide glycosylation selectively maintains the properties of cancer stem cells (CSCs) which is a serious clinical issue^16^. In addition, cancer cells overexpress thioredoxin (Trx) and deactivate ASK1 cell death pathway^17^ resulting in immune response suppression^18^. Tumors with low Trx levels exhibit a better prognosis than tumors with high Trx levels (poorer prognosis, *P* < 0.001) for partial free survival (PFS) and for overall survival (OS)^19^. GlcCer and Trx are the two biomarkers of resistance and prognostic recurrence^20^ respectively. Hence, tumor sensitizers targeting GlcCer and Trx which act as immunoadjuvants are presently the unmet medical need in cancer therapy.

**Scheme 1.**
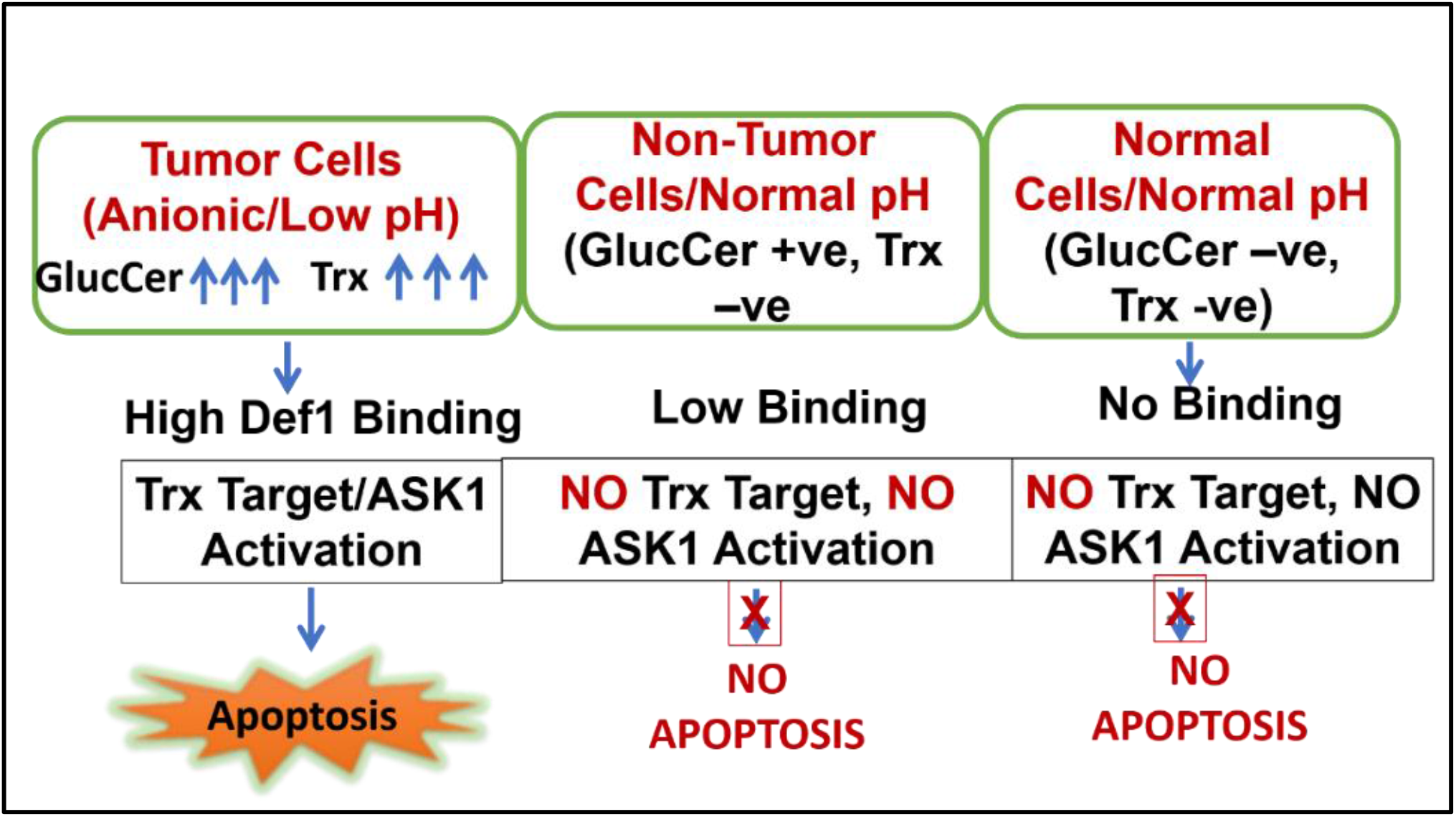
Def1 Selectivity

*Medicago sativa* Defensin 1 (MsDef1) is a natural antifungal peptide^21^ consisting of 45 amino acids folded with four disulfide bonds. MsDef1 is overexpressed in genetically modified potato to ward off a Verticillium wilt disease caused by a fungal pathogen *Verticillium dahlia* ^22^. Shah et.al.^23^ previously reported that a knockout of the *Fusarium graminearum* glucosylceramide synthase gene, Fg*gcs1*, blocked the antifungal activity of MsDef1 revealing the involvement of GlcCer in the mechanism of biological activity of this peptide. To date, no plant defensins have been shown to bind to MDR cells and synergize with chemotherapies. None of the current MDR modulators (e.g., Eliglustat) target two tumor specific MDR targets. Here, we report our data on MsDef1 as a targeted tumor sensitizer neoadjuvant to Doxorubicin. We show that MsDef1 targets two tumor specific targets i.e., membrane surface bound GlcCer and tumor intracellular Trx.

## Results

### 1. Synthesis of Def1

Def1 is a medium size tetra sulfide folded peptide with 45 amino acids which was synthesized by two methods; 1. recombination technology and 2. chemical synthesis followed by oxidation protocol. For recombinant method, Def1 which was extracted from the bacterial growth showed a number of bands in Western Blotting corresponding to different forms of Def1 (fully folded, partially folded etc., S1). The purification of the mixture by RP-HPLC yielded one pure fully folded Def1 with a corresponding isotopic molecular weight 5483. Def1 was also synthesized using chemical synthesis as an alternate method. First, linear peptide was synthesized using peptide synthesizer followed by a controlled air oxidation for 48 hrs to get fully folded Def1 corresponding to mono isotopic mass 5183). The kinetics of folding can be monitored through LC/MS/MS which gives the molecular ion peak on a real time basis. The final product was obtained by passing the solution through Sep-Pak and slow evaporation of the solvent get semicrystalline powder (Fig 1). It is to be noted that the oxidation protocol developed by us can be easily scaled up due to less impurities compared to recombinant method (see S1).

**Fig 1:**
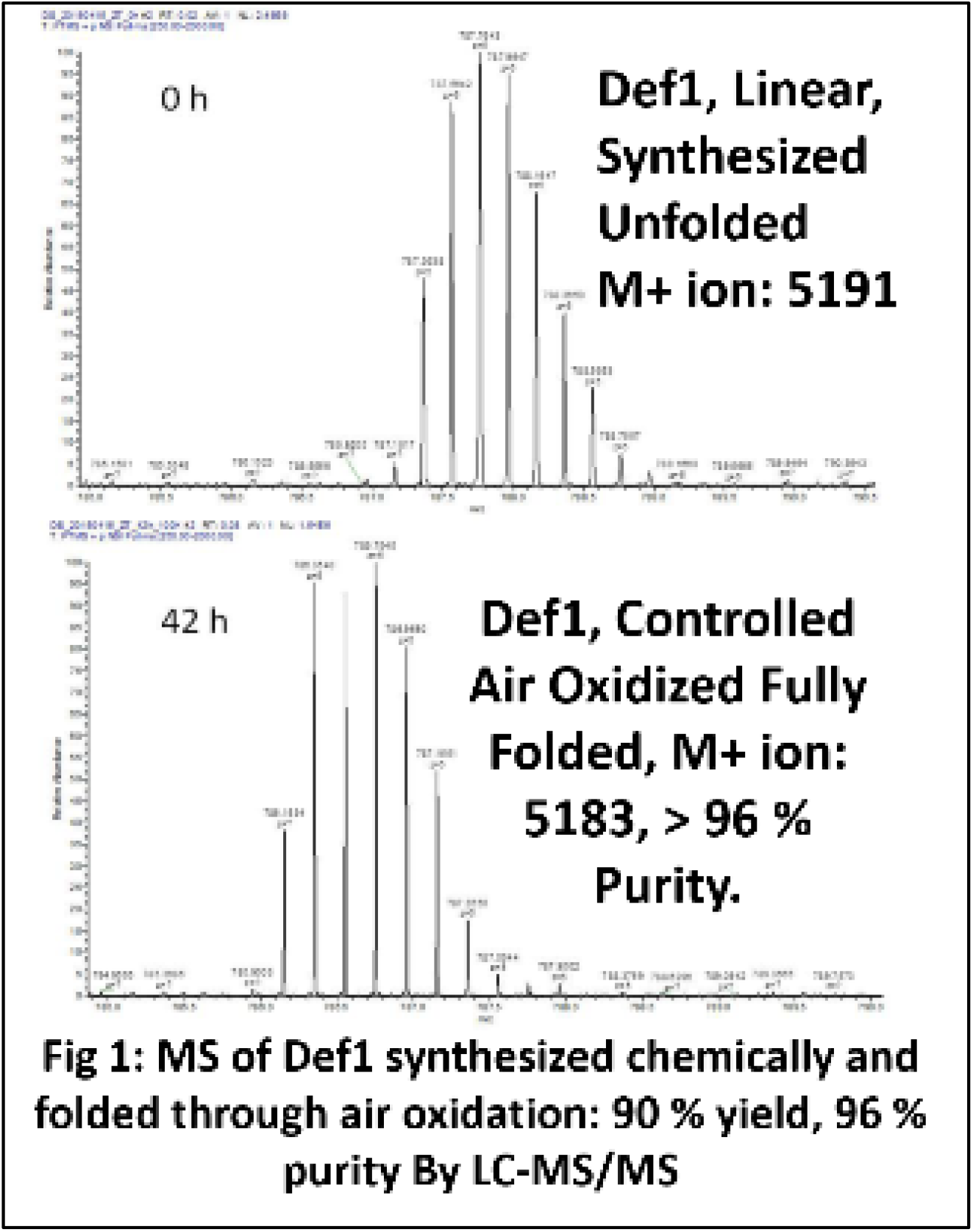
Characterization of Def1 using LC-MS and RP-HPLC.

### 2.A) Def1 binds to GlcCer “In Situ”

The solution dynamics of Def1 with GlcCer was assessed using chemical shift perturbation, ^15^N longitudinal relaxation (T1) and ^15^N-^1^H Nuclear Overhauser Effect (NOE). Def1 was labeled with ^15^N (NMR active nucleus) using the recombinant method where the active ingredient ^14^NH_4_Cl is replaced with ^15^N labeled NH_4_Cl. Our results *revealed that Def1 binds to GlcCer at two regions:* Asp36-Cys39 and amino acids between 12-20 and 33-40 (Fig. 2A and red portion of loop in B and C). In particular, we demonstrated that Arg38 is critical to binding since when Arg38 was mutated, it resulted in the complete loss of both binding and biological activity of Def1^24^. Similarly, linear Def1 did not show any biological activity emphasizing the role of tetra sulfide folding which helps to stabilize Def1 *in vivo* similar to many polysulfides. These results are compatible with the reported Psd1 defensin which showed Nuclear Overhauser Effect (NOE) with carbohydrate and ceramide parts of GlcCer^25^. This is the first premise on which Def1 is proposed as a targeted tumor sensitizer.

**Fig. 2:**
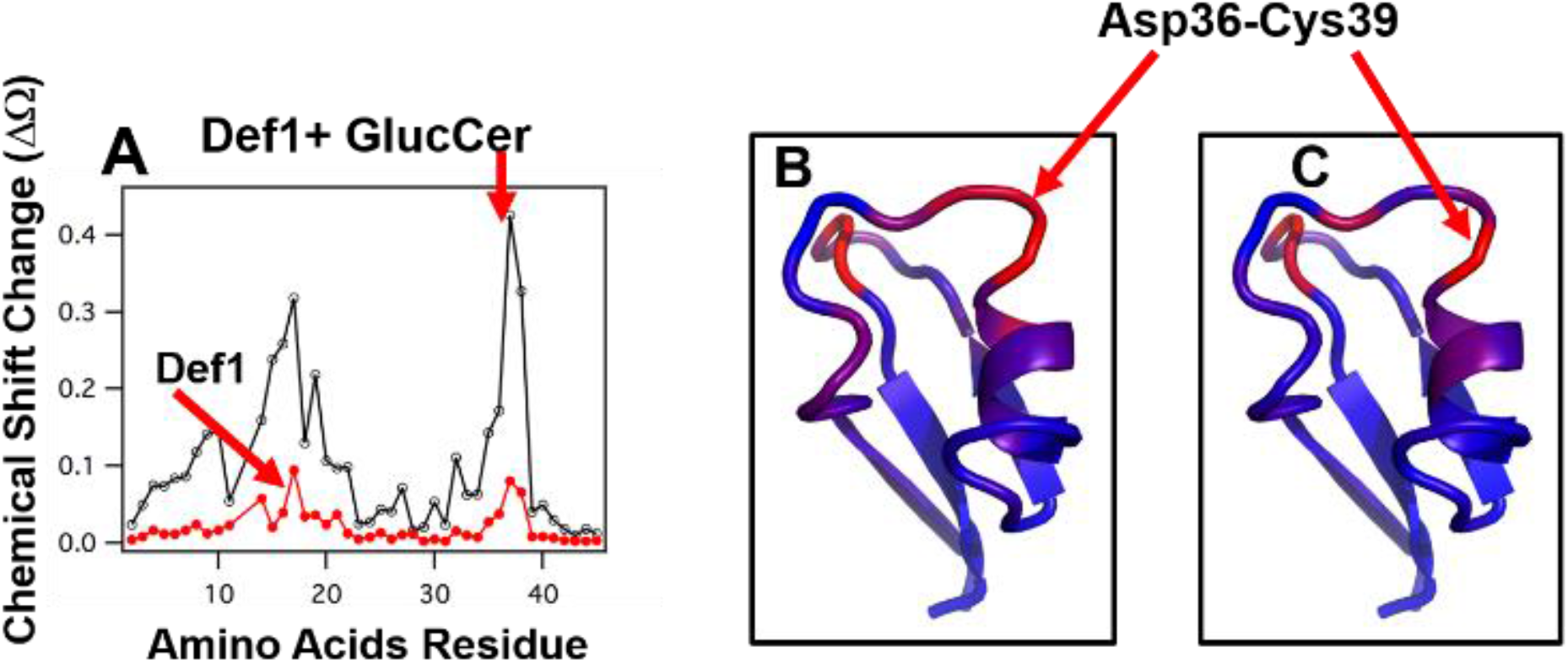
Binding of ^15^N-Def1 with Gluycosyl Ceramide (GlcCer) Using Chemical Shift Mapping by Solution ^15^N-NMR. A) Two areas of Def1 residues (Region 1: 12-20 and Region 2: 34-40 Residues) were bound to GlcCer. B-C: Two conformations of Def1 bound to GlucCer. The color scale extends from blue (no change, Δd=0) to red (significant chemical shift change, Δd≥0.25), N= 3, p <0.03.

### B). Def1 Regenerates Ceramide from GlcCer

Revamping ceramide pathway is important for those drugs which mediate cell death through ceramide pathway (e.g., Doxorubicin). Changes in the liberation of ceramide in Doxorubicin resistant MCF-7R cells were measured at Lipidomics Shared Resource, Medical University of South Carolina (MUSC), using high performance liquid chromatography-mass spectrometry (LC-MS/MS) as previously described by Jacek et.al.^26^, Preliminary studies (Fig 3) showed an enhanced accumulation of ceramide after 6 hrs of Def1 treatment (20 μM) in resistant breast cancer cells similar to positive control α-tocopheryl succinate (TOS, 20 μM)^27^ or compared to normal control cells (MCF-10A). Reactivation of ceramide pathway is important since it is an effective sensitizing strategy in overcoming the resistance in metastatic colon and breast cancers *in vivo*^28^. This is the second premise on which Def1 is proposed as a targeted tumor sensitizer.

**Fig 3:**
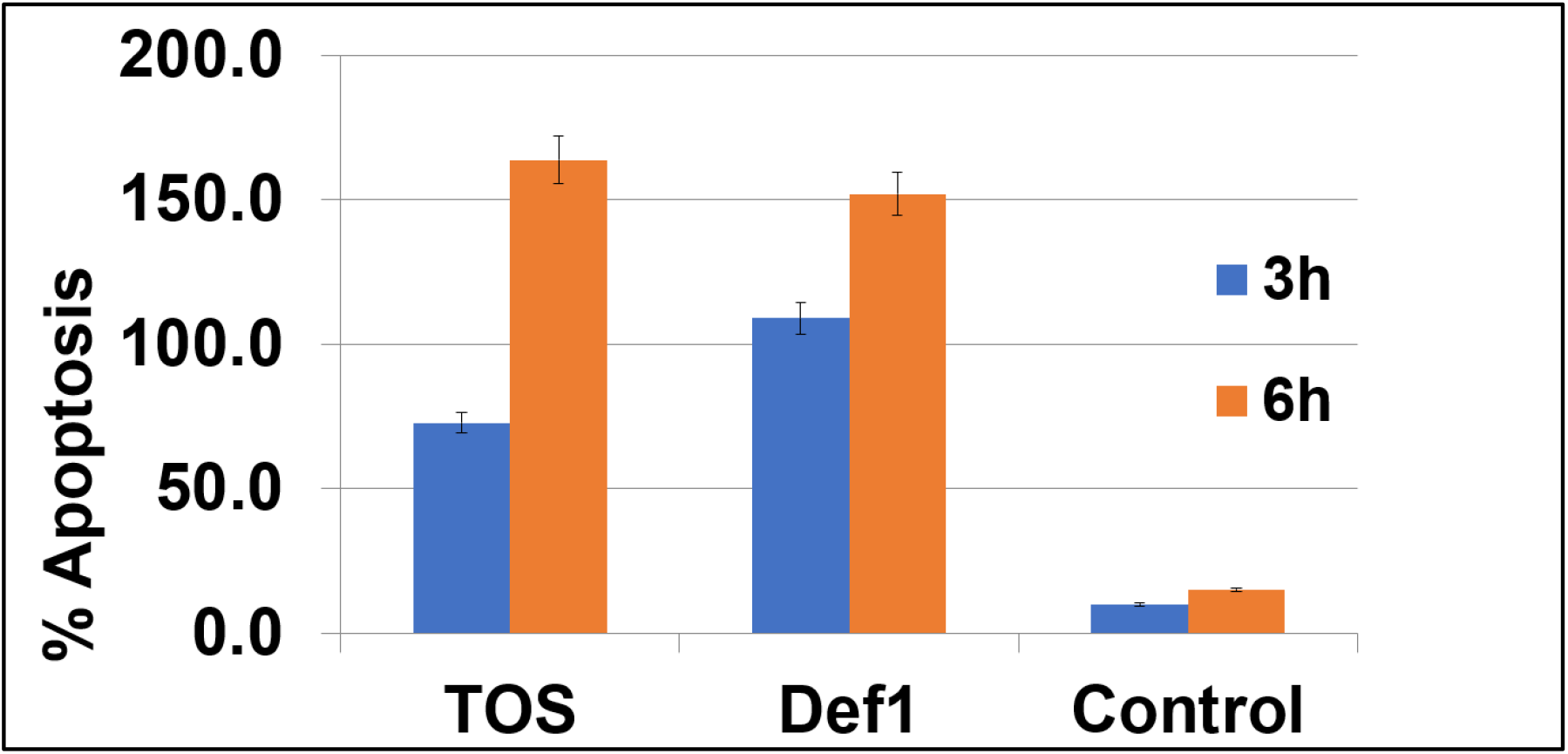
Def1 releases ceramide from GluCer on MDR MCF-7R cells (3 & 6hrs), Positive Control: a-Tocopheryl Succinate (TOS) and Control: MCF-10A (GlucCer negative, all 20 mM), n =4, p < 0.04.

### C). Def1 Oxidizes Tumor Specific Biomarker Thioredoxin (Trx)

Several tetra sulfide peptide derivatives are shown to oxidize thioredoxin (Trx) which is a tumor specific biomarker. Trx is known to involve in creating resistant cells by deactivating cell death pathway apoptosis stimulating kinase 1 (ASK1). Def1 was anticipated to interact with Trx due to four S-S bonds and hence, further studies of Def1 were conducted in cancer cells (e.g., triple negative breast cancer cells TNBC MDA-MB-231). Doxorubicin resistant MDA-MB-231-R cells (> 1.5 million) were treated with 0, 10 and 20 μM Def1 and 2 mM H_2_O_2_ (positive control). Fig 4a clearly shows oxidation of Trx at 20 μM Def1 by showing a new band at 28 KDa compared to untreated control which showed Trx band at 14 KDa. Oxidation of Trx by Def1 matches the positive control 2 mM H_2_O_2_ (N=4, p < 0.05). This is the third premise on which Def1 is proposed as a targeted tumor sensitizer.

**Fig 4.**
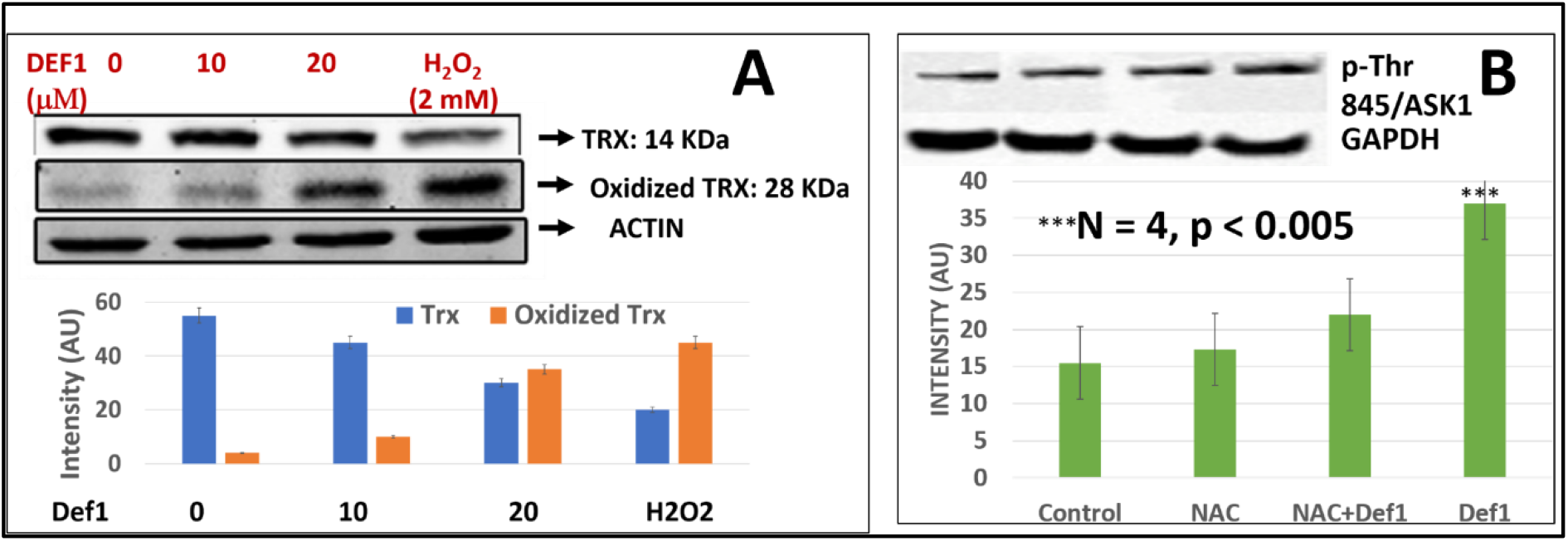
A: Def1 oxidizes Trx at 20 mM dose compared to positive control H_2_O_2_ (2 mM) & presumably activates ASK1 cell death pathway in MDA-MB-231-R TNBC cells, N=4, p < 0.05, B: : Immunoblot analysis of phosphorylation of Threonine-845 residue of ASK1 Protein in response to the treatment of Def1 in MDA-MB-231 cells: 1. Control, 2. N-Acetyl-Cysteine (NAC, 5 mM)/60 mins, 3. NAC (5 mM) + Def1 (50 μM) for 60 min, 4. Def-1 (50 μM) alone for 60 minutes. GAPDH: Internal Control.

### D) Def1 Disrupts Trx-ASK1 Complex to Activate ASK1 Cell Death Pathway and Induces Apoptosis in Resistant Cancer Cells

Oxidation of Trx is known to release ASK1 from the complex through phosphorylation^19b^. We corroborated Trx oxidation data (Fig 4a) by assessing the phosphorylation status of Thr845 residue of ASK1 using phosphorylation specific antibody Kit (Santa Cruz Biotech, CA). Fas resistant triple negative MDA-MB-231 cancer cells were treated with a) 50 μM Def1 for 60 minutes, b) N-acetyl cysteine (NAC, 5 mM) a negative control and c) NAC pretreated cells with Def1 (50 μM). Fig 4b showed a significant increase in phosphorylation of Thr845 residue (lane 4) compared to solvent control (lane 1) and NAC control (lane 2, N= 4, p < 0.005). NAC is FDA approved drug which inhibits phosphorylation of Thr-485 residues of ASK1 protecting cells from death. The higher intensity of phosphorylated residue (lane 4) for Def1 alone, <u>despite the additional inhibitive effect of NAC</u> indicates that Def1 targets tumor specific Trx and activates ASK1 cell death pathway. Trx inhibitors are known to reactivate ASK1 pathway and sensitize cancer cells to chemotherapy^29a^.

### E). Def1 Permeates Resistant Cancer Cells

Defensin type of molecules are cationic, particularly at tumor environmental low pH conditions which are known for <u>creating irreversible structural defects</u> on the cell membranes with a pore formation similar to cell penetrating peptides (CPPs)^40b^. However, the specificity of Def1 to MDR tumor cells need to be tested. Optical marker 6-((N-(7-nitrobenz-2-oxa-1, 3-diazol-4-yl) amino) hexanoyl (NBD) sphingosine was modified with linear Def1 before it was cyclized through oxidative protocol developed by us. The modified NBD-Def1 showed IC_50_ (12 μM) similar to the native Def1 indicating that modification of Def1 did not alter biological activity by a large margin (Table 1). Confocal microscopy studies on Def1-NBD which was incubated with resistant TNBC MDA-MB-231R and ovarian SKOV3 cells showed concentration dependent uptake of Def1-NBD (Fig 5A-B) compared to untreated tumor cells while, normal epithelial breast cells (MCF-10A) and fibroblasts cells (Fig 5C-D) did not pick up Def1 even at 5 times higher dose (200 mg/mL). Simple washing of stained cells did not reduce the fluorescence intensity originated from NBD which confirms trapping of Def1 inside the cancer cells. Permeation of membrane compromised fungal cells by Def1 was recently reported by us using optical marker DyLight 550-Def1^24b^. The low uptake of scrambled Def1-NBD by tumor cells (Fig 5E) establishes the selectivity of Def1 to tumor cells. The quantitation of Def1 uptake is shown in the graph (Fig 5F). Similarly, BODIPY-labeled NaD1, similar to Def1 is reported to have localized in organelles of lymphoma U937 cells^30a^.

**Table 1:**
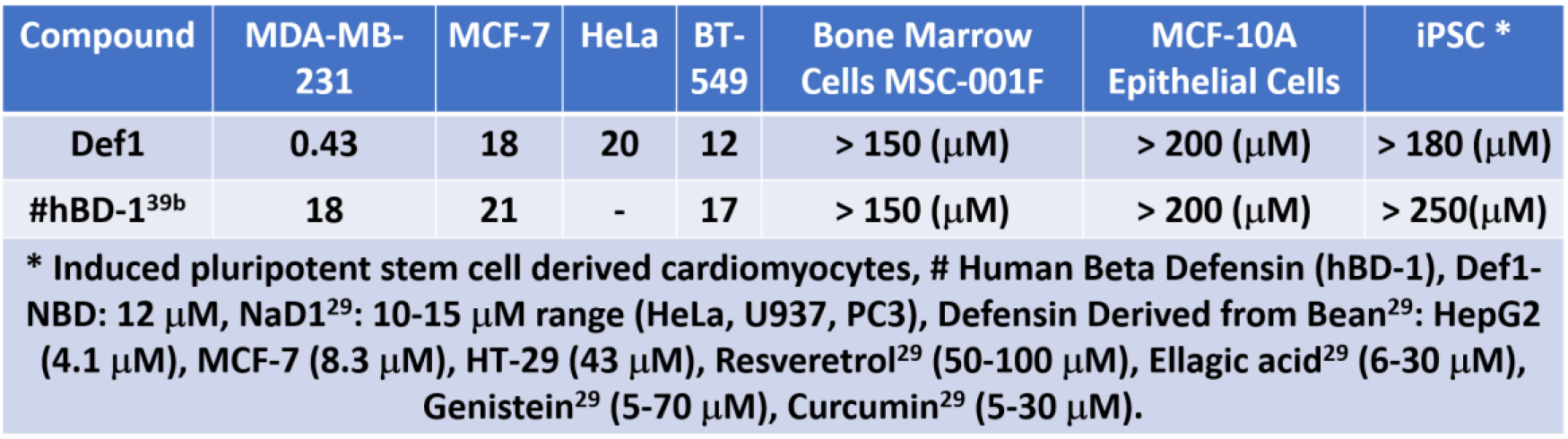
IC-50 of Plant Defensin (Def1) and Similar Defensins & Natural Tumor Sensitizers.

**Fig. 5:**
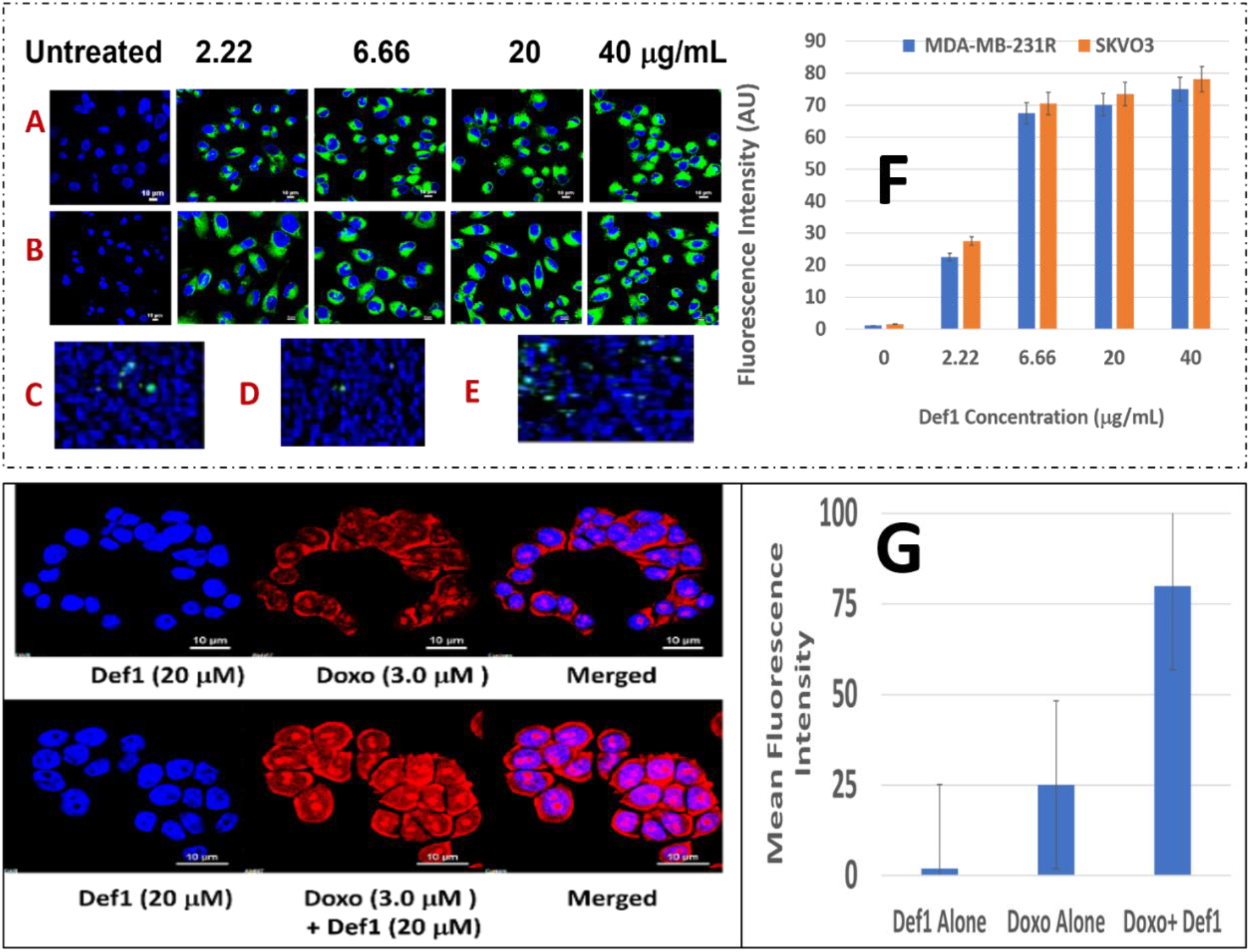
A: NBD-Def1 permeates resistant A) TNBC MDA-MB-231R Cells B) SKOV3 Cells while, uptake is low for C) Normal Epithelial Breast Cells D) Fibroblasts & E) Scrambled Def1 indicating selectivity, F) Graphic representation of Def1 uptake (N= 4, p < 001), G: Effect of Def1 on doxorubicin influx into MDR cancer cells MDA-MB-231R, N = 4, P < 0.002. Cells were stained with Hoechst 3325 and doxorubicin fluorescence was visualized at 488 nm.

### F). Def1 Enhances Diffusion of Def 1 in Resistant TNBC Cells

Def1 presumably compromises MDR cell membrane integrity by disintegrating GlucCer to ceramide and allowing Def1 influx to interact with the intracellular Trx. The hypothesis was corroborated by measuring the influx of Doxorubicin by Def1 in MDR cancer cells through the intrinsic fluorescence of Doxorubicin at 488 nm using confocal microscopy^10^. Fig 5G demonstrates the significant increase in the fluorescence intensity of Doxorubicin in MDA-MB-231R cells after Def1 treatment (20 μM) for 6-12 hrs compared to Doxorubicin alone. The increase in the fluorescence intensity correlated to Doxorubicin concentration is almost 3 times compared to neat Doxorubicin alone. This result resolves the poor tumor penetration limitation of Doxorubicin and may explain a) potential destruction of structure of cancer cell membranes and the synergistic effects of Def1 allowing Doxorubicin to perfuse better into the tumor.

### G). Def1 Shows Reasonable Antitumorigenic Activity

Table 1 shows IC-50 values for Def1 on several GlcCer and Trx positive cancer cells which are in the similar range as many chemotherapeutics^31^. Several similar defensin types of molecules derived from natural sources also showed IC-50 values in similar range^29-32^. However, none of them are shown to have multiple characteristics of Def1 including liberation of ceramide, oxidation of Trx and synergy between Def1 and chemotherapeutics. It is to be noted that Def1 seems to target cancer cells *in vitro* (e.g., MDA-MB-231, MDA-MB-231R, HeLa, BT-459) while sparing normal epithelial breast cells (e.g., MCF-10A), bone marrow cells (MSC-001F) and induced pluripotent stem cells derived cardiomyocytes (iPSC) which are important in determining the potential side effects of Def1. It is to be noted that Doxorubicin, the first line for TNBC treatment showed 9.6 μM IC-50 in iPSC cardiomyocytes compared to > 180 μM for Def1 indicating a better safety profile for Def1. Fig 6a shows the graphical representations of IC-50 values for cancer cells and normal cells. Fig 6B shows a representative IC-50 curves for the combination of Def1 at a fixed dose (20 μM) as a function of doxorubicin dose. There is a significant change in IC-50 combination (∼ 25 times) compared to either Def1 or doxorubicin. The synergy is verified through the generation of isobologram in Fig 6c.

**Fig 6:**
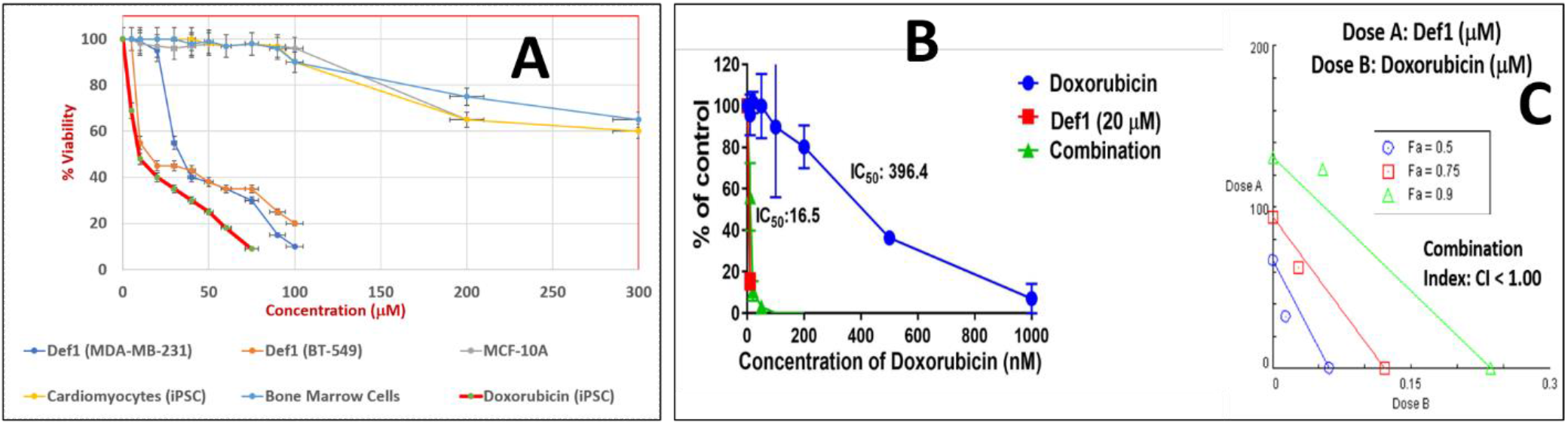
A: IC-50 values of Def1 in a) MDA-MB-231, b) BT-549 cancer cells and in normal cells c) MCF-10A, d) induced pluripotent stem cell derived cardiomyocytes, e) bone marrow cells (MSC-001F) and f) doxorubicin. The IC-50 values were derived from viability assays, N=3, p < 0.05, B: IC-50 curve for a) Def1 and b) combination of Def1 at 20 mM as a function of Doxorubicin, in MDA-MB-231 cells, N= 4, p< 0.002. C: Combination Index calculation for Def1 and doxorubicin in MDA-MB-231 cells: fa= fractional killing of cells. CI < 1.00 Synergistic, CI = 1.00 Additive, CI > 1.00, Antagonist.

### H) Def1 Synergizes with Doxorubicin

Although, Def1 is cytotoxic to cancer cells, its activation potential to sensitize low responsive MDR cancer cells will have a major impact on improving the clinical performance of chemotherapy. Synergy studies were conducted by varying both Def1 and Doxorubicin drugs to evaluate IC-50 values for the combination. Both Def1 and Doxorubicin showed a reasonable cell death (∼ 30 %) individually at low doses compared to untreated controls (Fig 7 A-C,). However, when the cancer cells were pretreated with 25 μM of Def1 followed by Doxorubicin treatment (0.7 μM), the combined treatment showed synergistic significant increase in cell death (> 75 % Fig 7D & 7H and graph) compared to either Doxorubicin or Def1 alone treatment (Fig. 7B-C or F-G). However, comparison of IC-50 values through viability assays showed that a combination of Def1 and Doxorubicin has IC-50 value ∼ 10 times lower compared to Doxorubicin (Fig 6B). Synergistic cell death was also verified by calculating the combination index (CI) values which were in the range 0.82-0.56 (for Def1 dose range 20-100 μM, Fig 10 B). CI was calculated using Chaou-Talalay method^41^ indicating that the combination is synergistic (CI < 1.00) and is not just additive (CI =1). CI values were obtained over a range of fractional cell kill levels (*i*.*e*., 0.05 to 0.95; 5-95% cell kill), and demonstrated values < 1 for two dose combinations of Def1 with Doxorubicin (Fig 6C, blue and red lines). This is the fourth premise on which Def1 is proposed as a targeted tumor sensitizer.

**Fig 7:**
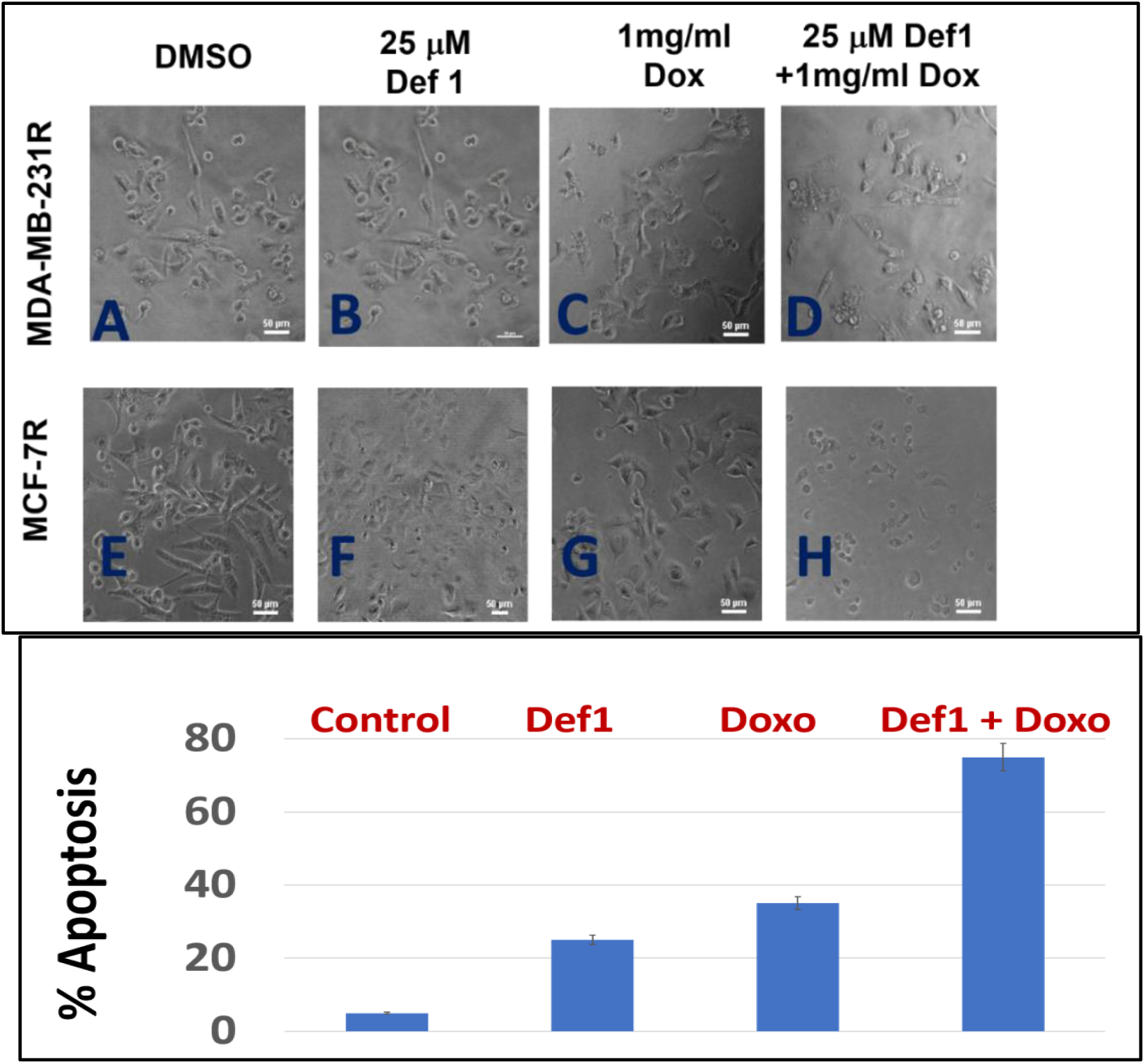
Def1 synergizes with doxorubicin in resistant MDA-MB-231R and MCF-7R cells A-D: Micro photographs of A & E) control, B & F) Def1 alone, C & G) Doxorubicin alone and D &H) Combination followed by quantification of apoptosis, N = 4, p < 0.007.

### I) Def1 Stability to Protease Digestion

The most important parameter that determines drug efficacy and safety is the stability of drug “*in vivo*”. The *in vivo* conditions are mimicked in our investigation by incubating Def1 with pepsin simulated gastric fluid (SGF). Def1, in presence of SGF showed little change in the intensity of the bands (lanes 3-6 compared to lanes 1-2 in Fig 8), while positive control BSA (lanes 9-1 in Fig 8) has degraded. This implies high stability of Def1 towards protease digestion. The combination of a cyclic cysteine knots (CCK) and a circular backbone renders peptide impervious to enzymatic breakdown exemplified by cyclotides^33^, theta defensins^34^ and FDA approved drugs like AcuTect^35^ and NeoTect^35^ developed by Pandurangi et.al^35^.

**Fig. 8:**
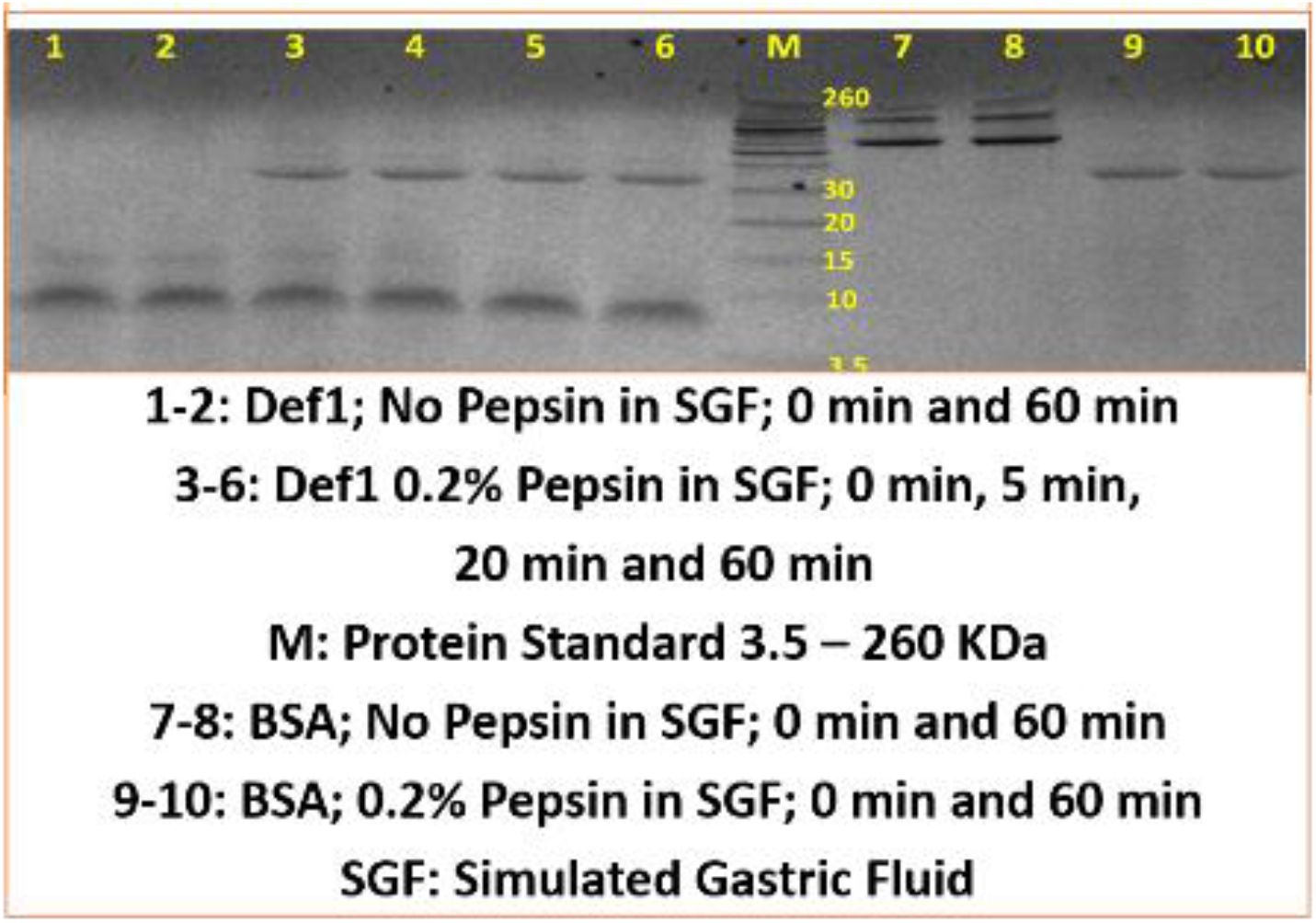
Def1 stability in Simulated Gastric Fluid (SGF, *In Vivo* mimicking conditions) while, positive control BSA degraded, n=4, p <0.08.

## Discussion

### Synthesis and Rationality for Extending Def1 Antifungal agent to Anticancer Agent

MsDef1 (Medicago Sativa Def1) is a natural protein available in food chain (alfalfa)^44^ with a specific sequence of 45 amino acids folded through four S-S bonds. Def1 is also overexpressed in corn and potato to ward off fungal pathogen^22^. The antifungal properties of Def1 were well established by Shah et.al^21-23^. particularly in Fusarium graminearum fungus. Def1 has a highly conserved a-core and c-core motifs determined by the homology-based models (using the I-TASSER website) with S-S folding (see supplementary materials S1).

Extensive research into the mechanism of action by Def1 on F. graminearum indicates a strong interaction between GlcCer on the surface of fungus and a cationic Def1. Apparently, the membrane vulnerability of fungi and cancer cells seem to have few common characteristics which enabled us to predict and propose the potential utility of Def1 in cancer therapeutics. For example, membrane fluidity and phospholipid content of fungal cells and cancer cells are similar. Def1, being cationic permeates fungal cells and cancer cells which are acidic and hence, makes Def1 more cationic (from +4 to +7) due to the protonation of amino acids in Def1. Higher the cationicity, better the permeation power as demonstrated by many cell permeable peptides (CPPs^40^). The major common factor between fungal cells and cancer cells for the current investigation is triggered from the GlcCer component of the membrane surface in them.

The major problem in the current cancer treatments is the creation of multi-drug cancer (MDR) cells by the treatment itself. For example, Doxorubicin which is a front-line treatment for triple negative breast cancer (TNBC) patients mediates cell death through the ceramide pathway. However, cancer cells circumvent the benefits of the therapy by glycosylating ceramide using glucosylceramide synthase (GCS) enzyme converting lethal ceramide to innocuous glucosylceramide (GlcCer)^16^. GlcCer is established as a biomarker for MDR^14^. Hence, targeting GlcCer may be a novel and innovative strategy for addressing clinical MDR issues.

Despite many cancer treatments are available, chemotherapy is still the first line of treatment for cancer. Chemotherapy is a non-specific therapy which suffers from off-target dose related acute and chronic side effects including creation of new drug resistant cells. For triple negative breast cancer (TNBC), the majority of patients are still treated with chemotherapy combination, despite the recent FDA approval of new PARP inhibitors therapies for TNBC^36^. While TNBC occurs in only 10–15% of all breast cancer patients, it accounts for almost half of all breast cancer deaths and the prognosis of metastatic TNBC is poor. Both nonspecific and targeted chemotherapy treatments can leave behind cancer resistant cells (CRCs)^4^, which remain the major cause of therapeutic failure in cancer treatments. This is a serious clinical problem which results in an increase in recurrence rate (13% for kidney cancer, 36% for breast and almost 100% for brain cancer) and make tumors refractory to future treatments^6^. CRCs, as a distinct population cause relapse and metastasis for greater than 40% of breast cancer patients, inducing <u>chemoresistance</u>^36^. Cancer cells desensitize themselves to intervention by activating survival pathways (e.g., ceramide and ASK1)^37^. None of the standard therapies, as far as we know are designed to address the molecular biology of the desensitization process and targets GlcCer which is established as MDR biomarker. Hence, Def1 as a targeting agent to MDR cancer may be feasible through GlcCer binding.

Our studies on the binding of Def1 with GlcCer using ^15^N NMR in solution identifies specific residues of Def1 which are important in causing the biological activity. MDR cancer cells (e.g., MDA-MB-231R) are characterized through the overexpression of GlcCer on their surface. Def1 has a positive charge 4, while in acidic conditions it changes to +7. The high anionic content of cancer cell membranes characterized through externalization of phosphatidyl serine facilitates high binding of cationic Def1. GlcCer was extracted from Doxorubicin resistant MCF-7R tumor cells as described by our collaborator Jacek et.al^26^. The solution dynamics of Def1 with GlucCer was assessed using chemical shift perturbation, ^15^N longitudinal relaxation (T1) and ^15^N-^1^H Nuclear Overhauser Effect (NOE). Def1 was labeled with ^15^N (NMR active nucleus) using the recombinant method where the active ingredient ^14^NH_4_Cl is replaced with ^15^N labeled NH_4_Cl. Our results revealed that Def1 binds to GlcCer at two regions: a) Asp36-Cys39 and b) amino acids between 12-20 and 33-40 (Fig. 2A and red portion of loop in B and C). In particular, we demonstrated that Arg38 is critical to binding since when Arg38 was mutated, it resulted in the complete loss of both binding and biological activity of Def1^8^. Similarly, linear Def1 did not show any biological activity emphasizing the role of tetra sulfide folding which helps Def1 to stabilize *in vivo* similar to many theta defensins^34^. These results are compatible with the reported Psd1 defensin which showed Nuclear Overhauser Effect (NOE) with carbohydrate and ceramide parts of GlcCer^25^. Binding of Def1 with GucCer is the first step in mediating the cytotoxicity and is the first scientific premise on which we rationalized Def1 for a potential cancer therapy.

### B). Def1 and Cell Death Pathways

Ceramide pathway is the most common way for many anthracyclines mode of action. Our results on resistant TNBC cells showed the release of ceramide from GlcCer (Fig 3). This is exactly opposite to cancer cells tactics to nullify the cytotoxic effects of ceramide by glycosylating ability of cancer cells to convert apoptotic ceramide to non-apoptotic GlcCer. In other words, Def1 treatment may revamp the ceramide pathway inhibiting one of the resistance mechanisms. Our data showed that Def1 induced sphingomyelinase activity and enhanced ceramide levels in resistant TNBC cells (Fig 3). The importance of restoration of ceramide pathway lies in its ability to modulate the biochemical and cellular processes that will lead to apoptosis. Def1 was also compared with the positive control alfa Tocopheryl succinate (α-TOS) which mediates its cytotoxic effect through ceramide pathway^27^. The significant increase in the ceramide release by Def1 in resistant cancer cells may suggest that Def1 presumably may be able to create structural defects on the surface of cancer cell membranes by cleaving the bond between ceramide and glucose.

Equally important in the development of MDR is the overexpression of thioredoxin as a defense mechanism in response to stimuli including oxidative stress mediated through chemotherapy by deactivating cell death pathway ASK1. High Trx levels are directly correlated to inhibition of endogenous apoptosis stimulating kinase 1 pathway making chemotherapy ineffective. Tumors with low Trx levels exhibited a better prognosis than tumors with high Trx levels (poorer prognosis, *P* < 0.001) for partial free survival (PFS) and for overall survival (OS)^19^. In fact, Trx is a well-studied target which lowers the response rate to specific docetaxel, cis-platin and Doxorubicin. Oxidation of Trx releases ASK1 bonded to Trx. Fig 3 clearly shows oxidation of Trx at 20 μM Def1 compared to 2 mM H_2_O_2_ which is a positive control making Def1 to possess a better oxidation potential than H_2_O_2_. Oxidation of Trx is known to activate ASK1 cell death pathway and sensitizes tumor cells^38^. Disintegrating GlcCer positive MDR cancer cells by Def1 may be rationalized through the access of tumor intracellular Trx. This result corroborates well with human beta defensin (hBD-1) which oxidizes Trx *in situ* (see Fig 2 in ref. 39, HPLC data).

We corroborated Trx oxidation data by assessing the phosphorylation status of Thr845 residue of ASK1 using phosphorylation specific antibody Kit (Santa Cruz Biotech, CA). Fas resistant triple negative MDA-MB-231 cancer cells were treated with a) 50 μM Def1 for 60 minutes, b) N-acetyl cysteine (NAC, 5 mM) a negative control and c) NAC pretreated cells with Def1 (50 μM). Fig 4b showed a significant increase in phosphorylation of Thr845 residue (lane 4) compared to solvent control (lane 1) and NAC control (lane 2, N= 4, p < 0.005). NAC is FDA approved drug which inhibits phosphorylation of Thr-485 residues of ASK1 protecting cells from death. The higher intensity of phosphorylated residue (lane 4) for Def1 alone, <u>despite the</u> additional inhibitive effect of NAC indicates that Def1 targets tumor specific Trx and activates ASK1 cell death pathway. Trx inhibitors are known to reactivate ASK1 pathway and sensitize cancer cells to chemotherapy^38^. Targeting dual cell death pathways (ceramide and ASK1) is the hall mark of Def1 which may be unique compared to the other existing cancer treatments.

### C). Cell Penetrating Potential of Def1

Interaction of Def1 with the tumor specific MDR biomarker Trx may have several implications on the potential trapping of Def1 which may be essential for the continuous activation of ASK1 cell death pathway resulting in the antitumor properties of Def1 and/or synergy with the existing treatments. Defensin type of molecules are cationic at low pH which are known for cell penetrating properties (e.g., human beta defensin hBD-3^39^) with a pore formation capacity similar to cell penetrating peptides (CPPs)^40^. However, the specificity and cell penetrating ability of Def1 in MDR tumor cells need to be tested. Optical marker 6-((N-(7-nitrobenz-2-oxa-1, 3-diazol-4-yl) amino) hexanoyl sphingosine (NBD) was conjugated with linear Def1 before it was cyclized through oxidative protocol developed by us. The modified NBD-Def1 showed IC_50_ (12 μM, see Table 1) similar to Def1 indicating that modification of Def1 did not affect biological activity by a large margin (Table 1). Confocal microscopy studies on Def1-NBD incubated with TNBC MDA-MB-231R and ovarian SKOV3 cells showed concentration dependent uptake of Def1-NBD compared to untreated tumor cells (Fig 5A-B) while, normal fibroblasts and epithelial breast cells (Fig 5C-D) did not pick up Def1 even at 5 times higher dose (200 mg/mL). Simple washing of stained cells did not reduce the intensity originated from NBD-Def1 and confirms trapping of Def1 inside the cancer cells. Permeation of membrane compromised fungal cells by Def1 was recently reported by us using optical marker DyLight 550-Def1^24^. The low uptake of scrambled Def1-NBD by tumor cells (Fig 5E) establishes the selectivity of Def1 to tumor cells. BODIPY-labeled NaD1, similar to Def1 is reported to localize in organelles of lymphoma U937 cells^30^ demonstrating the cell penetrating ability of Def1.

Current treatments for TNBC patients revolve around chemotherapy (particularly, anthracyclines including Doxorubicin) despite non-specific and high off-target toxicity. The poor tumor penetrating power of Doxorubicin demands the administration of high dose which in turn induces more resistance over a period of time and also effluxed by the pumps leading to the low dose delivery to the inner parts of tumor. Def1 was shown to create a pore effects on the GlcCer positive cancer cell membranes which may, potentially increase the diffusion of drug into the tumor. The hypothesis was corroborated by measuring the influx of Doxorubicin by Def1 in MDR cancer cells through the intrinsic fluorescence of Doxorubicin at 488 nm using confocal microscopy. Fig 5b demonstrates the significant increase in the fluorescence intensity of Doxorubicin in MDA-MB-231R cells after Def1 treatment (20 μM). This result resolves the poor tumor penetration limitation of Doxorubicin^40^ and may explain the cumulative effects of Def1 induced cell death with higher dose of Doxorubicin.

### D). Def1 as a Potential Antitumor Agent

Table 1 shows IC-50 values for Def1 in several GlcCer and Trx positive cancer cells which are in the similar range as many chemotherapeutics^31^. Notably, normal cells including human induced pluripotent stem cell derived cardiomyocytes (iPSCs) and bone marrow cells have 5-7 times higher IC-50 values compared to cancer cells which indicates a reasonable selectivity of Def1 for cancer cells. In contrast, the standard chemotherapy Doxorubicin shows 10-15 times more potency for both cardiomyocytes and bone marrow cells which resulted in off-target toxicity *in vivo*. However, it is interesting to know that when Doxorubicin was combined with Def1, the IC-50 of the combination is ∼ 25 times less in MDA-MB-231 cells which indicates a potential synergy between Def1 and Doxorubicin (Fig 9A). The data was corroborated with the calculation of combination index (CI) using Towley method^41^. Different combination of two drug doses were titrated to assess fractional kills at 0.5, 0.75 and 0.9 fa equivalent to 50, 75 and 90 % cell death to generate isobologram. The synergy is defined for those drug dose combinations which are below the line, while the point on the line is additive and the point above is antagonistic^41^. Fig 9 B shows the drug doses correspond to blue and redlines are synergistic while for green line drug dose combination it is antagonistic. Although, several defensins^32^ have been reported to be cytotoxic to cancer cells they have different homology and none of them were shown to bind GluCer, permeates cells, synergize with Doxorubicin (see combination index, CI values < 1, Table 2) and hit the intracellular tumor specific target (Trx). This is the **fourth premise** on which Def1 is proposed as a targeted tumor sensitizer.

### E). Def1 and Synergy with Chemotherapy

The risk of anthracycline related cardiomyopathy increases with a higher cumulative anthracycline dose^42^: About 3% of that dose persist even after the completion of therapy leading to potential heart failure and death in 400 mg/m^2^, 7% for a dose of 550 mg/m^2^, and 18% for a dose of 700 mg/m^2^. The poor tumor penetration by Doxorubicin and off targeted hyperactivation of endogenous PARP in heart leads to cardiomyopathy^43^. Hence, improving the clinical performance of Doxorubicin is an unmet medical need. A priori activation of apoptosis pathways of tumor (AAAPT) technology developed by us involves targeted natural tumor sensitizers to sensitize specifically desensitized resistant tumor cells to evoke a better response from Doxorubicin^24c^. This synergistic approach may facilitate lower use of Doxorubicin by increasing the therapeutic index of the drug. Both Def1 (20 μM) and Doxorubicin (10 μM) induced cell death in resistant TNBC MDA-MB-231-R cells reasonably assessed through morphology of cell death (Fig 10). However, when combined together, the cumulative cell death was greater than just adding the two drugs. The change in IC-50 for the combination is ∼ 10 times higher compared to individual drugs. This translates, clinically that a combined formulation potentially may reduce the tumor burden at a lower Doxorubicin dose which in turn may reduce the dose related toxicity due to Doxorubicin. In other words, physicians may have a larger window for fine tuning the dose regimen based on the combined toxicity rather than individual toxicity. Def1, being natural and commercialized in genetically modified corn is presumably not toxic at the clinical dose levels.

Two major requirements for the translation of bench concept to clinical product are a) stability of Def1 and b) selectivity of Def1 to cancer cells. In general, Def1 type of molecules are quite stable in media due to the cyclization of cysteines. Stability of Def1 *in vivo* towards proteolytic enzymes may not be a big concern since several tetra sulfide array compounds including FDA approved products developed by Pandurangi et.al; (e.g., NeoTect, NeoTide), human beta defensin (hBD-1) with three S-S bonds, cyclized cyclotides^33^, theta defensins^34^ provides solid examples of high stable compounds similar to Def1. Typical concentration of defensin involved in host defense against microbial infections range ∼ 10mg/mL (e.g., granules, intestine) indicates reasonable stability *in vivo*^*44*^. Preliminary studies on the stability involved incubation of Def1 in simulated gastric fluid (SGF) which mimics *in vivo* conditions, and the stability was assessed through Western Blotting. Def1 showed no significant change in the intensity of the bands with respect to time, while positive control BSA was degraded easily (Fig 11). This implies high stability of Def1 towards protease digestion. The combination of a cyclic cysteine knots (CCK) and a circular backbone renders peptide impervious to enzymatic breakdown exemplified by cyclotides^33^, theta defensins^34^ and FDA approved drugs like AcuTect^35^ and NeoTect^35^.

### F). Def1 Selectivity

Def1 is shown to target two tumor specific targets in this present study, i.e., GlcCer and Trx. Overexpression of both GlcCer and Trx made cancer cells resistant to treatments and are considered as biomarkers of resistance. The selectivity of Def1 may be hypothesized due to <u>the combination</u> of a) it’s high binding to MDR specific GlcCer, b) oxidation of the Trx and c) higher cationicity of Def1 (7+) at low pH tumor environment (Scheme 1) which interact with anionic membranes of cancer cells. The non-target cells may have either GlcCer positive and Trx negative or cells with both GlcCer and Trx negative. Low binding of Def1 to cells is presumably does not release therapeutic dose of ceramide or enough Def1 dose reaches intracellular target Trx to modify it. Undesired off-target expression of GlcCer and Trx (e.g., brain, kidney etc.) may not be a major concern. It is to be noted that overexpression of GlcCer and Trx in tumor is the key. It is the relative target Vs. non-target expression of the biomarker which makes it better selective compared to nonspecific chemotherapy. High target/non-target ratio for many FDA approved targeted imaging and therapeutic drugs (e.g., Octreoscan^45^, Herceptin^46^ based on somatostatin and ErbB2 overexpression respectively, NeoTect developed by us and PX-12 (Trx inhibitor^29^) are good examples of selectivity despite low expression of the biomarker at non-target sites. It is the combination of two tumor specific targets (GlcCer & Trx) which are involved in the desensitization process and Def1 reactivated the two corresponding cell death pathways (ceramide and ASK1) which resulted in sensitizing MDR cancer cells (Scheme 2).

**Scheme 2.**
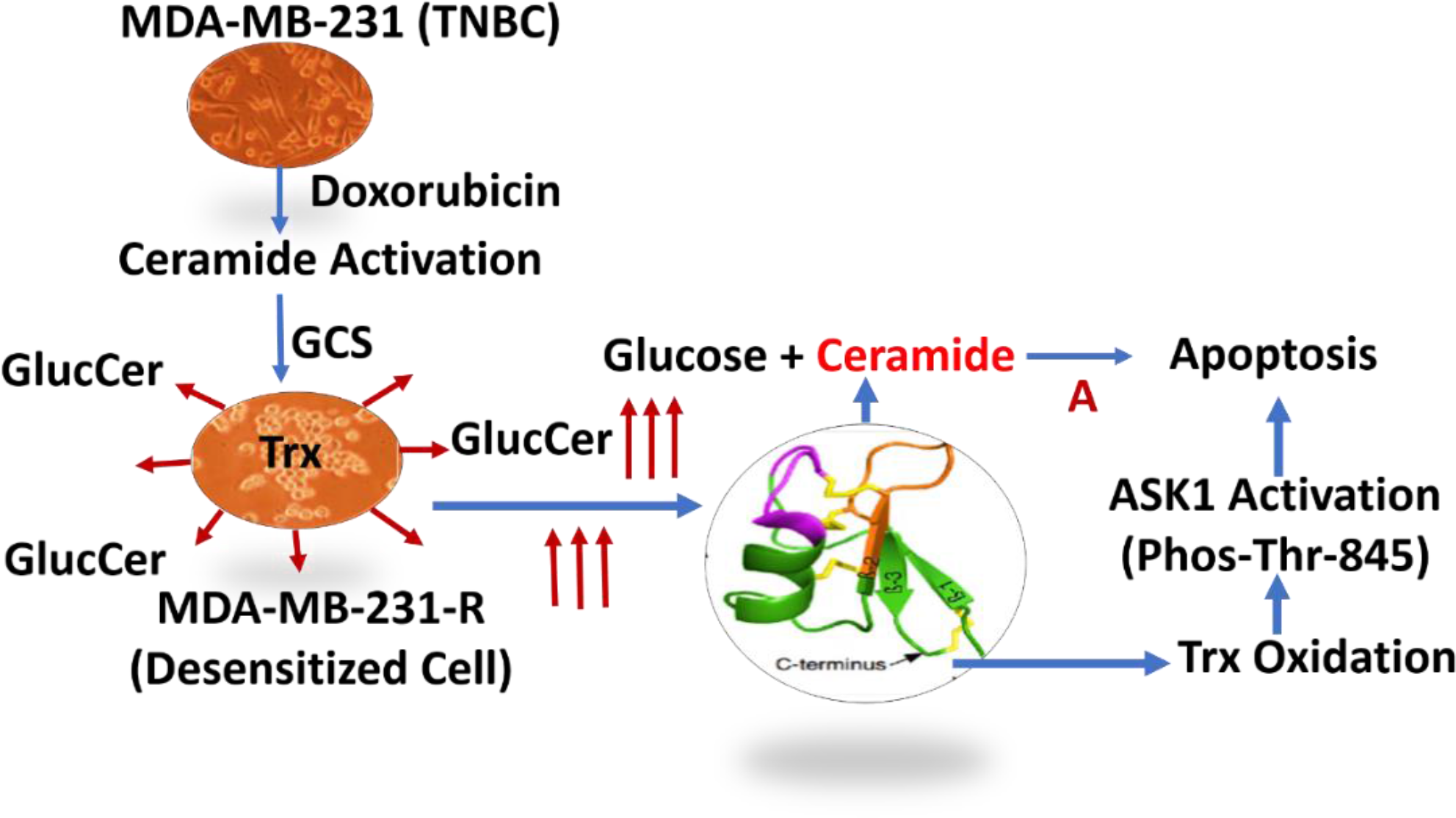
Def1 Pathways to Sensitive MDR Cancer Cells

In summary, TNBC patients treated with Doxorubicin are not getting benefitted much since ceramide pathway through which Doxorubicin mediates cell death was deactivated by cancer cells. The problem is due to the classic glycosylation of ceramide to GlcCer (Scheme 2) producing MDR cancer cells. Def1 reactivates two specific cell death pathways which presumably sensitized MDR cells resulting in the synergistic effects with Doxorubicin. Studies revealed that Def1 could be a natural tumor sensitizer which can be useful at neoadjuvant settings with chemotherapy. Further studies are needed to test the synergicity of Def1 in tumor animal models for a potential smooth clinical translation.

## Materials and Methods

### 1. Synthesis of MsDef1

MsDef1 was produced by using two methods; 1) recombinant expression in *P. pastoris* as described previously^23^ 2) controlled air oxidation of the linear peptide synthesized by using standard peptide synthesizer (Apex 396 Parallel Synthesizer). Def1 prepared by recombination method was further purified by RP-HPLC column chromatography using a reverse phase C18 column (Delta Pak Wat 011793, 15063.9 mm, 5 mM, 300 A) to obtain 95% purity and characterized by mass spectroscopy. MsDef1 was dissolved in sterile double distilled water and its concentration was determined by the BCA assay kit (Pierce, Rockford, IL). MsDef1 concentration was also determined by quantitative amino acid analysis performed at the Proteomics and Mass Spectrophotometry. MsDef1 was filtered through 0.22 µm filter before using it for *in vitro* antifungal activity assays. A solution of linear Def1 was dissolved in double distilled water in a test tube fitted with a probe through which air is bubbled through the solution for 36-48 hrs and monitored through mass spectrometry for a molecular ion corresponding to oxidation of four S-S bonds. For example, 0.4 mg of the sample was dissolved to 0.6 μg/μM in 50 mM phosphate buffer containing 1M Gu HCl at pH 7.5. The sample was aliquoted at different time points: 0, 4h, 18h, 24h, 36 and 48h. The samples were desalted with C18 zip tip and run on LTQ-Orbitrap Velos by direct infusion. The samples were run with high resolution (60,000, LTQ-Velos Pro Orbitrap LC-MS/MS). Analytical: Def1: HPLC > 96 5 Pure, Mol. Wt.: Linear: (M+H) =5195.1, (deconvoluted mono isotopic mass = 5191.23), Folded: (M+H) = 5187.25, (deconvoluted mono isotopic mass = 5183.23, Fig 1 shows the characterization of Def1 by slow oxidation method which coincides with recombinant method (see S1 for impurity profile of Def1 before HPLC purification).

### 2. Synthesis of

6-((N-(7-nitrobenz-2-oxa-1, 3-diazol-4-yl) amino) hexanoyl (NBD)-Def1: The crude linear Def1 was dissolved in 20% methanol/8 M guanidinium hydrochloride and mixed with 6-((N-(7-nitrobenz-2-oxa-1, 3-diazol-4-yl) amino) hexanoyl (NBD) sphingosine in dark conditions. The mixture was stirred for 48 hrs before passing through the sep-pak and purified with RP-HPLC using eluents of H_2_O/0.1% TFA (eluent A) and acetonitrile/0.1% TFA (eluent B). The programmed elution profile 0–1 min, 100% A; 1–80 min, B is increased from 0–75% at a flow rate of 10 mL/min on preparative column (Spirit Peptide C18, 5 µm column, 25 × 2.12) Peptide purity was determined by analytical HPLC monitoring peptide elution by absorbance at 220 nm. The peptide mass was characterized by matrix assisted laser desorption/ionization (MALDI) on a Voyager DE Pro (Applied Biosystems Inc., Foster City, CA). The pure Def1 conjugate was air oxidized slowly with continuous stirring for 36-48 hrs and monitored the formation of S-S folding using MS/MS, Molecular (M+H) = 5745.9 corresponding to the fully folded NBD-Def1 conjugate.

### 3. Def1 Binding to GlcCer using ^15^N NMR

Synthesis of ^15^N labeled Def1: In order to study the solution dynamics of Def1 with GlcCer by NMR, Def1 was labeled with NMR active nucleus 15N. NMR experiments were conducted on a Varian 700MHz spectrometer with HCN probe (backbone dynamics) and Bruker 600 MHz spectrometer with QCI cryoprobe (assignments and structure determination). Backbone (N and H_N_) and side chain protons have been assigned for Def1 in aqueous buffer (25 mM HEPES, 20 mM NaCl, pH 6) at 25 °C. The backbone amides were assigned using ^15^N-separated TOCSY and NOESY spectra with 60 ms and 150 ms mixing times, respectively. These experiments were acquired at 25 °C with a 250 µM ^15^N-labeled Def1 sample in 90% H2O/10% D2O. Side chain protons were assigned using these spectra plus ^1^H-^1^H COSY, TOCSY, and NOESY spectra acquired for an otherwise identical unlabeled Def1 sample in 100% D2O. Backbone amide dynamics experiments were carried out using the same ^15^N-labeled Def1 sample. R_1_, R_2_, and heteronuclear NOE experiments were performed using standard pulse sequences. At least 8 timepoints were acquired for R_1_ and R_2_ measurements and heteronuclear NOEs were determined from an average of two independent experiments. Error estimates were based on the signal/noise ratio of each spectrum. Single exponential decays were fit to determine R_1_ and R_2_ rates using IgoPro (Wave metrics). Model free analysis was performed using fast MF and Model Free 4.15. Chemical shift changes upon interaction with lipid were determined using the ^15^N-labeled Def1 sample. 60 μM d25-DPC (dodecyl phosphocholine, per deuterated acyl tail) was added to the 0.25 μM Def1 sample in aqueous buffer to investigate the interaction of Def1 with DPC micelles. This sample was then added to lyophilized glucosylceramide, allowed to equilibrate for 2 hours, and returned to the NMR tube to look for additional changes reflecting specific interaction with glucosylceramide.

### 3. ^15^N Def1 Synthesis

Phosphatidylcholine (PC) was purchased from Aldrich. ^15^NH_4_Cl was purchased from Cambridge Isotope Laboratory. MsDef1 was expressed in Pichia pastoris and purified as described previously^23^. Briefly, Pichia cultures were grown overnight in buffered YNB media with no amino acids and 25 g. ^15^NH_4_Cl dissolved in 1M Potassium Phosphate (pH = 6.0), 500x Biotin, 10 % glycerol and then induced with methanol every 24 h, according to the manufacturer’s directions (Invitrogen). The cultures were grown for 7 d at 29°C, and cells were removed by centrifugation at 2,000g for 15 min. Defensin was purified from the growth medium by first dialyzing against 25 mM sodium acetate, pH 4.5, and then passing the dialysate through CM-Sephadex C-25 cation-exchange resin (Amersham Biosciences, Piscataway, NJ) equilibrated with 25 mM sodium acetate, pH 4.5. Resin was extensively washed with binding buffer (25 mM sodium acetate, pH 4.5), and the bound protein was then eluted in 1 M NaCl, 50 mM Tris, pH 7.6. Fractions containing the protein were manually collected and analyzed by SDS-PAGE for the presence of the defensin. Fractions containing the defensin protein were concentrated using a Minitan II ultrafiltration system (Millipore, Bedford, MA) with a 3-kD cutoff membrane and dialyzed against 10 mM Tris, pH 7.6. Purity was assessed using Homogenous 20 SDS gels on a Phastgel system (Amersham Biosciences) followed by reverse phase HPLC. The partially purified MsDef1 was further purified by reverse-phase HPLC column chromatography (Beckman Coulter, Brea, CA) using a reverse phase C18 column (Delta Pak Wat 011793, 15063.9 mm, 5 mM, 300 A) to obtain > 95% purity and characterized using MS. The typical MS shows a single peak at 5254.03 (M+H) corresponding to fully folded and labeled ^15^N-MsDef1.

### 4. Isolation of GlcCer from F. graminearum

About 1g of F.graminearum was powdered in nitrogen liquid followed by extraction with 2:1 methanol: chloroform (9 mL), centrifuged at 2500 rpm for 5 minutes and the supernatant was mixed with 3 mL chloroform and added water to a ratio 1:1: 0.9. The solution was centrifuged for 10 minutes before separating upper phase and lower phase which contains nonpolar phase to which few drops of ethanol was added to get the clear liquid, dried under stream of nitrogen slowly.

### 5. Purification of GlcCer

The nonpolar fraction was resuspended in 3 mL of 100:1 chloroform: acetic acid. A silica column (SPE DSC-Si Silica tube 6 ml (Supelco) Cat # 52655 –U pack) was equilibrated with 10 ml of 100:1 chloroform: acetic acid before the lipid was applied to the column. The silica column was washed with 10 mL of 100:1 chloroform: acetic acid before adding 5 mL of 80: 20 ratios of chloroform: acetone. Fractions in the next few steps were collected with (a) 10 mL of acetone (b) 8 ml of 100:1 acetone: acetic acid before they were mixed and dried using stream of nitrogen gas. The dried substance was suspended in 2 mL of chloroform, added 2 ml of 0.6 N methanolic sodium hydroxide, mixed well and kept at room temperature for 1 hr. The solution was acidified using 1.3 mL 1N HCl and 0.5 mL of water before centrifuged for 5 minutes. The lower phase is separated in a new tube and few drops of ethanol was added to dissolve completely and dried again. The dried substance was resuspended in 3 ml of 100:1 chloroform: acetic acid before putting on the silica column and eluted with (a) 5 mL 80:20 chloroform: acetone (b) 6 mL 50:50 chloroform: acetone twice each time 3mL. The column was washed with 8 mL acetone, 8 ml of 100:1 acetone: acetic acid, collected and dried. The next and final step was the hydrolysis of sphingolipids when the dried substance was resuspended in 2 ml of 2 N methanolic HCl and heated at 70°C for 8 h. after cooling the tube, 1mL of water was added, centrifuged and the upper phase was separated, dried, added heptanes and was analyzed by GC-MS. The GlcCer was further purified using RP-HPLC collecting all the fractions (1 mL) and each fraction was eluted on TLC (specification) to compare with the standard plant glucocerebroside (Matreya LLC, PA). The typical mass spectrum showed a single peak (> 95 % purity) at M+ ion 754.

### 6. Viability Assays and IC-50

Cell viability was evaluated by the MTT method. All cancer cells (MDA-MB-231, MCF-7, Hela, MCF-10A epithelial breast cells, induced pluripotent stem cells derived cardiomyocytes (iPSc) were plated in 96-well flat-bottom plates at the density of 2×10^4^ per well, allowed to attach overnight, and treated with different concentrations of Def1 (1-100 µM) After 24 h, 10 µl (3–4,5-dimethylthiazol-2-yl)-2,5-diphenyltetrazolium bromide (MTT; Sigma) were added to each well and the plate incubated at 37°C for 3 h. After removing the media, 200 µl of isopropanol were added to dissolve the crystals. Absorbance was read at 550 nm in an ELISA plate reader (Sunrise, Tecan, Milan, Italy), and the results are expressed as relative change with respect to the controls set as 100%. Human induced pluripotent stem cell cardiomyocytes (iPSCs) were used (Ionic Transports, St Louis) for assessing viability.

### 7. Thioredoxin Assays Using Western Blotting

The thioredoxin Western blots were performed as described previously, with minor modifications^29^. Approximately 2.5 million cells were lysed in G-lysis buffer (50 mM Tris–HCl, pH 8.3, 3 mM EDTA, 6 M guanidine–HCl, 0.5% Triton X-100) containing 50 mM iodoacetic acid (IAA; pH 8.3). For each experiment, control plates, for identifying thioredoxin redox state bands in the Western blot, were also incubated with 2mM H2O2, for 10 min at room temperature, before incubation with 50 mM IAA. Afterward, the lysate from all cells was incubated in the dark for 30 min, with the IAA. The lysates were then centrifuged in G-25 micro spin columns (GE Healthcare). Protein was then quantified from the eluent using the Bradford protein assay, as previously described earlier^24^. Equal amounts of protein were then added to a 15% acrylamide native gel. The gels were run at 100 V constant for approximately 2.5 h. The proteins contained in the gel were then transferred to a nitrocellulose membrane (Bio-Rad Laboratories) using a semidry transfer protocol. The nitrocellulose membrane was then washed in PBST (phosphate-buffered saline with 0.1% Tween) and blocked in 5% milk with PBST for 1 h, before being incubated at 4 °C overnight with the primary antibody, 1:1000 goat anti-hTrx-1 (American Diagnostica) in PBST with 2% bovine serum albumin. The primary antibody was then removed; the blot was washed in PBST for 10 min three times, with constant shaking, before being incubated for 1 h with the secondary antibody [horseradish peroxidase (HRP)-labeled rabbit anti-goat IgG (Santa Cruz Biotechnology, Cat. No. sc2020)]. The blot was then washed again for 10 min three times in PBST before being treated with HRP chemiluminescence detection reagents (Renaissance; NEN). The protein was then visualized by exposing the blot to X-ray film for 2–5 min in a dark room with a film cassette before developing the film. In an alternate method of protein visualization, a Typhoon FLA 7000 (GE Healthcare Life Sciences) imaging system was used to directly detect the fluorescent signal from the blotting.

### 8.0 Ceramide Assessment

Drug resistant MCF-7R cells were grown on 10 cm plates containing 2–5 X10^6^ cells per plate, were washed with 10 mL of PBS. The cells were scraped with 1 mL of methanol and transferred into a 5.5 mL glass vial with either aluminum or Teflon-sealed caps (SKS Science). Cells were sonicated for 60 min in a bath sonicator, and 100 μL of the solution was taken for protein concentration measurements. 50 μL of a 50 ng/mL C17 standard solution was added to the remaining cells in the glass vial to obtain a final concentration of 10 ng/ mL of C17 as the internal standard (IS). At the end of the extraction procedure, 2 mL of chloroform was added to the cell suspension. After vortexing for 5 s followed by 30 min of sonication, the cell lysates were spun for 5 min at 3000 rpm. Then, the lower layer containing chloroform were transferred to a new tube using a 1 mL glass pipette, leaving the upper layer (containing methanol) and the middle layers (containing proteins) (∼75% extraction efficiency for this first extraction). The tube with the layer containing chloroform and lipids was placed on ice. 2 mL of chloroform was added to the other two remaining layers for second extraction. The second extraction solution was combined with the first extraction solution (∼ 92% extraction efficiency for two extractions combined). The extraction efficiency was evaluated by a similar procedure to recovery experiments in which ceramide concentration was analyzed before and after extraction. The percentage of the measured concentration of ceramide after extraction over the measured concentration before extraction is the extraction efficiency. The extraction solution was dried under nitrogen gas at room temperature. 250 mL of acetonitrile was added to the completely dried vial and vortexed well. The acetonitrile reconstituted extraction solutions with lipids were stored in a -80 °C freezer until analysis. High-performance liquid chromatography (HPLC) was performed on an Agilent 1200 (Agilent Technologies, Santa Clara, CA) series instrument equipped with a quaternary pump and linked to an HTC PAL autosampler (CTC Analytics) with a compartment for six 96-well plates. A Pursuit 3 Diphenyl reversed-phase column, 50 × 2.0 mm, 3 mm (Varian, Walnut Creek, CA) was used, and the flow rate was set to 0.8 mL/min. A five-minute gradient was run for HPLC. Mobile phase A consisted of 0.1% formic acid and 25 mM ammonium acetate in water and mobile phase B consisted of 100% acetonitrile.

### 9.0 Permeability Assays

Resistant triple negative breast cancer cells MDA-MB-231R and ovarian SKOV3 cells were incubated with 20ug/ml, 6.6ug/ml or 2.2ug/ml of NBD-Def1 for 15 min @37C. (total volume = 200ul of media), washed off the excess NBD-Def1 with 1xPBS and fixed cells with 4% parafolmadehyde to be imaged using Nikon Confocal microscope. In order to determine the specificity of NBD-Def1 internalization, 1mg/mL neat Def1 was incubated and washed before imaging the cells. For assessing Doxorubicin influx in resistant MDA-MB-231 TNBC cells, auto fluorescence of Doxorubicin at signal emission at 595 nm post excitation with a 470 nm laser.

### 10. Def1 Stability

The Simulated Gastric Fluid (GSF) assay was performed as described by Fu et al., 2002 with some modification. Tests were performed in 500 μL of SGF (200 mg NaCl, 0.2% pepsin, pH 2) in glass tubes in a 37°C water bath with continuous stirring of the enzyme reaction. After 2 min preincubation, the assay was started by the addition of 25 μL of MsDef1 peptide (5 mg/ml) to each vial containing SGF, SGF without pepsin or ultrapure water. Bovine serum albumin (minimum 98%, A-7030, Sigma Aldrich) (5 mg/ml in ultrapure water) was used as the positive control for pepsin digestion. Protein samples were analyzed for degradation by SDS PAGE followed by staining for 1 h and extensive washing with ultrapure water.

## Acknowledgements

Part of this research was supported by the grant SBIR NIH R43CA183353 and R43CA214223 Funding organization did not play any roles in designing experiments, data collection, analysis or any decision to publish the data. NIH has provided only financial support to Sci-Engi-Medco Solutions Inc (SEMCO) through salary (RP, PI) and research cost.

## Captions for Supporting Information

S1: A: SDS-PAGE chromatography of Def1 prepared using recombinant method, B; HPLC peaks for the gel showing several peaks.

## Supplementary Materials

**S1: Fig 1:**
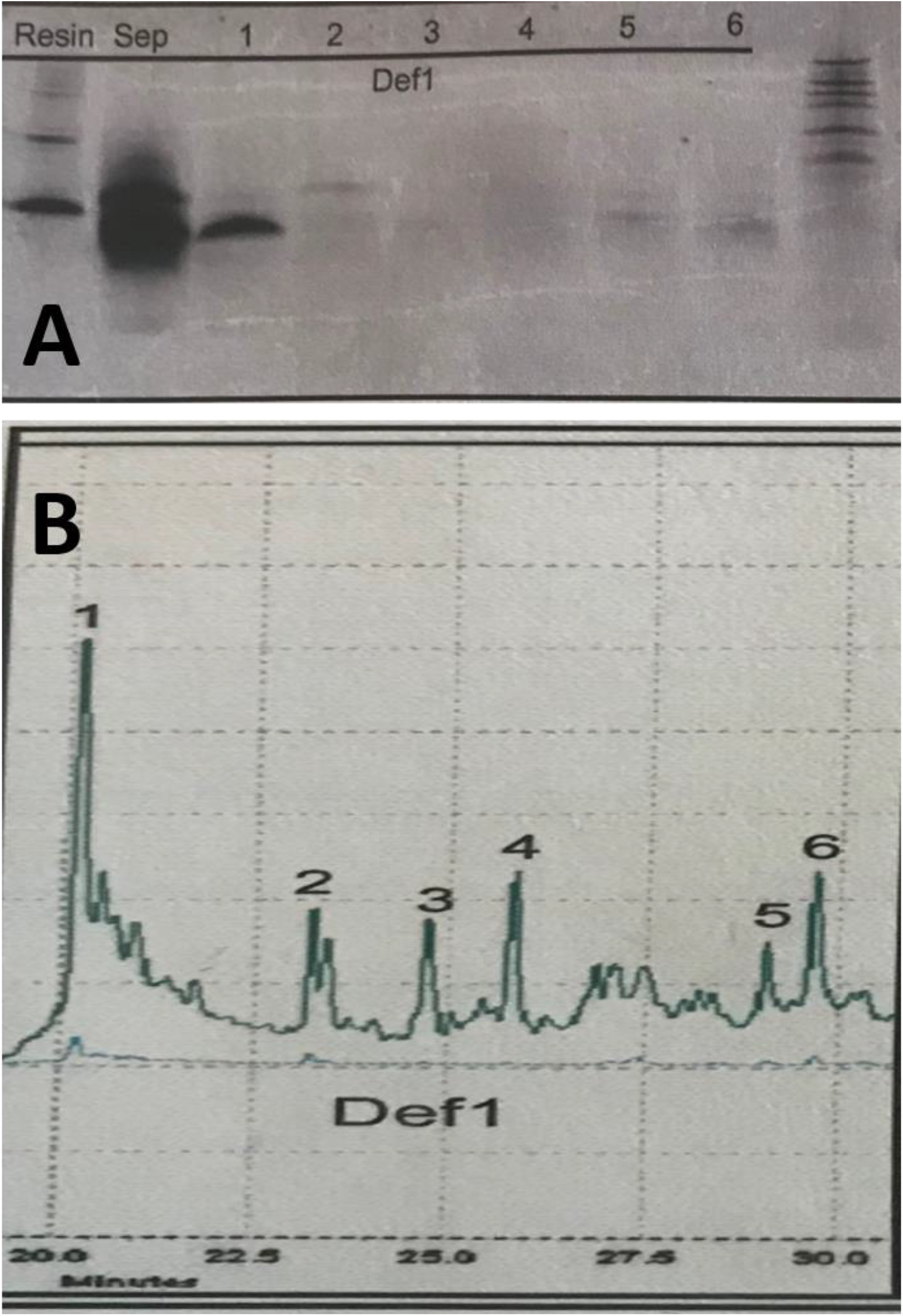
A: SDS-PAGE of Def1 by recombinant method using E.Coli Rosetta (DE3)/pET28a, Line 1: Protein eluted from Talon resin, Line 2: Protein eluted from SepPac (after His-tag cleavage), 1-6: HPLC fractions, Standard reference marker. B: HPLC chromatogram for Def1.

## Bibliography

1. a) Szakács G, Paterson JK, Ludwig JA, Booth-Genthe C, Gottesman MM. Targeting multidrug resistance in cancer. Nat Rev Drug Discov. 2006 Mar;5(3):219–34. doi: 10.1038/nrd1984. PMID: 16518375. b) Trédan, O., Galmarini, C. M., Patel, K., & Tannock, I. F. (2007). Trédan O, Galmarini CM, Patel K, Tannock IF. Drug resistance and the solid tumor microenvironment. J Natl Cancer Inst. 2007 Oct 3;99(19):1441–54. doi: 10.1093/jnci/djm135. Epub 2007 Sep 25. PMID: 17895480. c) Wu Q, Yang Z, Nie Y, Shi Y, Fan D. Multi-drug resistance in cancer chemotherapeutics: mechanisms and lab approaches. Cancer Lett. 2014 Jun 1;347(2):159–66. doi: 10.1016/j.canlet.2014.03.013. Epub 2014 Mar 19. PMID: 24657660.

2. Jin, Z.; El-Deiry, W. S. Overview of Cell Death Signaling Pathways. Cancer Biology and Therapy. 2005. https://doi.org/10.4161/cbt.4.2.1508.

3. a) Evans C, Dalgleish AG, Kumar D. Review article: immune suppression and colorectal cancer. Aliment Pharmacol Ther. 2006 Oct 15;24(8):1163–77. b) Markman JL, Shiao SL. Impact of the immune system and immunotherapy in colorectal cancer. J Gastrointest Oncol. 2015;6(2):208–223.

4. a) Regnard, C., Kindlen, M., Regnard, C., Kindlen, M. Chemotherapy: Side Effects. In Supportive and Palliative Care in Cancer; 2019, b) Monsuez, J. J., Charniot, J. C., Vignat, N., Artigou, J. Y. Cardiac Side-Effects of Cancer Chemotherapy. International Journal of Cardiology. 2010. c) Love, R. R., Leventhal, H., Easterling, D. V., Nerenz, D. R. Side Effects and Emotional Distress during Cancer Chemotherapy. Cancer 1989. Shapiro, C. L., Recht, A. Side Effects of Adjuvant Treatment of Breast Cancer. New England Journal of Medicine 2001. https://doi.org/10.1056/nejm200106283442607.

5. a) Kumar, P., Aggarwal, R. An Overview of Triple-Negative Breast Cancer. Archives of Gynecology and Obstetrics. 2016. https://doi.org/10.1007/s00404-015-3859-y, b) Bauer, K. R., Brown, M., Cress, R. D., Parise, C. A., Caggiano, V. Descriptive Analysis of Estrogen Receptor (ER)-Negative, Progesterone Receptor (PR)-Negative, and HER2-Negative Invasive Breast Cancer, the so-Called Triple-Negative Phenotype: A Population-Based Study from the California Cancer Registry. Cancer 2007. https://doi.org/10.1002/cncr.22618, c) Wahba, H. A., El-Hadaad, H. A. Current Approaches in Treatment of Triple-Negative Breast Cancer. Cancer Biology and Medicine. 2015. https://doi.org/10.7497/j.issn.2095-3941.2015.0030.

6. a) Reddy SM, Barcenas CH, Sinha AK, et al. Long-term survival outcomes of triple-receptor negative breast cancer survivors who are disease free at 5 years and relationship with low hormone receptor positivity. Br J Cancer. 2017;118(1):17–23.

7. Shelley, M., Harrison, C., Coles, B., Staffurth, J., Wilt, T. J., Mason, M. D. Chemotherapy for Hormone-Refractory Prostate Cancer. Cochrane Database of Systematic Reviews. 2006. https://doi.org/10.1002/14651858.CD005247.pub2.

8. Mi-Joung Lee et.al; Lymphedema Following Taxane-Based Chemotherapy in Women with Early Breast Cancer, Lymphatic Research and Biology 2014 Dec;12(4):282–8

9. a) Duda, D. G., Kozin, S. V., Kirkpatrick, N. D., Xu, L., Fukumura, D., Jain, R. K. CXCL12 (SDF1α)-CXCR4/CXCR7 Pathway Inhibition: An Emerging Sensitizer for Anticancer Therapies? Clinical Cancer Research 2011. https://doi.org/10.1158/1078-0432.CCR-10-2636. b) Kvols, L. K. Radiation Sensitizers: A Selective Review of Molecules Targeting DNA and Non-DNA Targets. Journal of Nuclear Medicine. 2005, Jan;46 Suppl 1:187S–90S.

10. a) Reddy SM, Barcenas CH, Sinha AK, et al. Long-term survival outcomes of triple-receptor negative breast cancer survivors who are disease free at 5 years and relationship with low hormone receptor positivity. Br J Cancer. 2017;118(1):17–23, b) Dietze EC, Sistrunk C, Miranda-Carboni G, O’Regan R, Seewaldt VL. Triple-negative breast cancer in African-American women: disparities versus biology. Nat Rev Cancer. 2015 Apr;15(4):248–54.

11. Siddharth, S. & Sharma, D. Racial Disparity and Triple-Negative Breast Cancer in African-American Women: A Multifaceted Affair between Obesity, Biology, and Socioeconomic Determinants. Cancers (Basel). 10, (2018), Dec 14;10(12):514. doi: 10.3390/cancers10120514

12. a) Evans C, Dalgleish AG, Kumar D. Review article: immune suppression and colorectal cancer. Aliment Pharmacol Ther. 2006 Oct 15;24(8):1163–77. b) Markman JL, Shiao SL. Impact of the immune system and immunotherapy in colorectal cancer. J Gastrointest Oncol. 2015;6(2):208–223.

13. Lavie Y, Cao H, Bursten SL, Giuliano AE, Cabot MC. Accumulation of glucosylceramides in multidrug-resistant cancer cells. J Biol Chem. 1996, 271(32):19530–6.

14. Margaret A. Park The relationship between multidrug resistance and glucosylceramide levels: An opportunity for combined therapies, Cancer Biology & Therapy, 8:12, 2009,1122–1124.

15. Kartal Yandım, M., Apohan, E. & Baran, Y. Therapeutic potential of targeting ceramide/glucosylceramide pathway in cancer. Cancer Chemother Pharmacol 71, 13–20 (2013). https://doi.org/10.1007/s00280-012-1984-x.

16. Gupta V, Bhinge KN, Hosain SB, Xiong K, Gu X, Shi R, Ho MY, Khoo KH, Li SC, Li YT, Ambudkar SV, Jazwinski SM, Liu YY. Ceramide glycosylation by glucosylceramide synthase selectively maintains the properties of breast cancer stem cells. J Biol Chem. 2012, 287(44):37195–205.

17. a) Grogan, T. M., Fenoglio-Prieser, C., Zeheb, R., Bellamy, W., Frutiger, Y., Vela, E., Stemmerman, G., Macdonald, J., Richter, L., Gallegos, A., Powis, G. Thioredoxin, a Putative Oncogene Product, Is Overexpressed in Gastric Carcinoma and Associated with Increased Proliferation and Increased Cell Survival. Human Pathology 2000. https://doi.org/10.1053/hp.2000.6546, b) Kaimul, A. M., Nakamura, H., Masutani, H., Yodoi, J. Thioredoxin and Thioredoxin-Binding Protein-2 in Cancer and Metabolic Syndrome. Free Radical Biology and Medicine. 2007. https://doi.org/10.1016/j.freeradbiomed.2007.05.032.

18. a) Michaud, M., Martins, I., Sukkurwala, A. Q., Adjemian, S., Ma, Y., Pellegatti, P., Shen, S., Kepp, O., Scoazec, M., Mignot, G., Rello-Varona, S., Tailler, M., Menger, L., Vacchelli, E., Galluzzi, L., Ghiringhelli, F., Di Virgilio, F., Zitvogel, L., Kroemer, G. Autophagy-Dependent Anticancer Immune Responses Induced by Chemotherapeutic Agents in Mice. Science 2011. https://doi.org/10.1126/science.1208347. b) Zitvogel, L., Apetoh, L., Ghiringhelli, F., Kroemer, G. Immunological Aspects of Cancer Chemotherapy. Nature Reviews Immunology. 2008. https://doi.org/10.1038/nri2216, c) Kam, T., Alexander, M. Drug-Induced Immune Thrombocytopenia. Journal of Pharmacy Practice. 2014. https://doi.org/10.1177/0897190014546099.

19. Woolston, C. M., Zhang, L., Storr, S. J., Al-Attar, A., Shehata, M., Ellis, I. O., Chan, S. Y., Martin, S. G. The Prognostic and Predictive Power of Redox Protein Expression for Anthracycline-Based Chemotherapy Response in Locally Advanced Breast Cancer. Modern Pathology. 2012. https://doi.org/10.1038/modpathol.2012.60.

20. a) Liu, Y. Y., Hill, R. A., Li, Y. T. Ceramide Glycosylation Catalyzed by Glucosylceramide Synthase and Cancer Drug Resistance. In Advances in Cancer Research; 2013. https://doi.org/10.1016/B978-0-12-394274-6.00003-0, b) Shan, W., Zhong, W., Zhao, R., Oberley, T. D. Thioredoxin 1 as a Subcellular Biomarker of Redox Imbalance in Human Prostate Cancer Progression. Free Radical Biology and Medicine 2010. https://doi.org/10.1016/j.freeradbiomed.2010.10.691.

21. Sathoff, A. E., Velivelli, S., Shah, D. M., Samac, D. A. Plant Defensin Peptides Have Antifungal and Antibacterial Activity against Human and Plant Pathogens. Phytopathology 2019. https://doi.org/10.1094/PHYTO-09-18-0331-R.

22. Shah, D. M., Snyder, A. K. Antifungal Plant Proteins and Generation of Transgenic Plants That Inhibit the Growth of Phytopathogenic Fungi., 2008, PCT applied.

23. Abdallah, N. A., Shah, D., Abbas, D., Madkour, M. Stable Integration and Expression of a Plant Defensin in Tomato Confers Resistance to Fusarium Wilt. GM crops 2010. https://doi.org/10.4161/gmcr.1.5.15091, b) Ramamoorthy, V., Cahoon, E. B., Li, J., Thokala, M., Minto, R. E., Shah, D. M. Glucosylceramide Synthase Is Essential for Alfalfa Defensin-Mediated Growth Inhibition but Not for Pathogenicity of Fusarium Graminearum. Molecular Microbiology 2007. https://doi.org/10.1111/j.1365-2958.2007.05955.x. b) Ramamoorthy, V.;Zhao, X., Snyder, A. K., Xu, J. R., Shah, D. M. Two Mitogen-Activated Protein Kinase Signalling Cascades Mediate Basal Resistance to Antifungal Plant Defensins in Fusarium Graminearum. Cellular Microbiology 2007. https://doi.org/10.1111/j.1462-5822.2006.00887.x.

24. a) Sagaram US, Pandurangi R, Kaur J, Smith TJ, Shah DM (2011) Structure-activity determinants in antifungal plant Defensins MsDef1 and MtDef4 with different modes of action against Fusarium graminearum. PLoS ONE 6(4): e18550, b) Sagaram US, El-Mounadi K, Buchko GW, Berg HR, Kaur J, Pandurangi RS, et al. (2013) Structural and Functional Studies of a Phosphatidic Acid-Binding Antifungal Plant Defensin MtDef4: Identification of an RGFRRR Motif Governing Fungal Cell Entry. PLoS ONE 8(12): e82485. https://doi.org/10.1371/journal.pone.0082485, c) AAAPT Technology: Raghu Pandurangi,Compositions and methods for sensitizing low-responsive tumors to cancer therapy, PCT Filed, Jan 2016, PCT/US16/68554, Publication number, WO/2017/131911, Publication date: 08/03/2017, International filing date: 12/23/2016, https://patentimages.storage.googleapis.com/6d/3a/2b/2e778b81566dbc/WO2017131911A1.pdf

25. a) De Medeiros LN, Domitrovic T, de Andrade PC, Faria J, Bergter EB, Weissmüller G, Kurtenbach E. Psd1 binding affinity toward fungal membrane components as assessed by SPR: The role of glucosylceramide in fungal recognition and entry, Biopolymers. 2014 Nov;102(6):456–64, b) Amaral VSGD, Santos SACS, de Andrade PC, et al. Pisum sativum Defensin 1 Eradicates Mouse Metastatic Lung Nodules from B16F10 Melanoma Cells. Int J Mol Sci. 2020;21(8):2662.

26. Jacek Bielawski, Jason S. Pierce, Justin Snider, Barbara Rembiesa, Zdzislaw M. Szulc, and Alicja Bielawsk, Gomprehensive Quantitatiue Analysis of Bioactiue Sphingolipids by High-Performance Liquid Ghromatography-Tandem Mass Spectrometry, Donald Armstrong (ed.l, Lipidomics, Methods in Molecular Biology, vol. 579, pp 22–58.

27. Xinbin Gu, Xiaodong Song, Yongheng Dong, et al. Vitamin E Succinate Induces Ceramide-Mediated Apoptosis in Head and Neck Squamous Cell Carcinoma In vitro and In vivo. Clin Cancer Res 2008, 14:1840–1848.

28. Chen, C. L., Lin, C. F., Chang, W. T., Huang, W. C., Teng, C. F., Lin, Y. S. Ceramide Induces P38 MAPK and JNK Activation through a Mechanism Involving a Thioredoxin-Interacting Protein-Mediated Pathway. Blood 2008. https://doi.org/10.1182/blood-2007-08-106336.

29. a) Ramanathan RK et.al; A randomized phase II study of PX-12, an inhibitor of thioredoxin in patients with advanced cancer of the pancreas following progression after a gemcitabine-containing combination. Cancer Chemother Pharmacol. 2011 Mar;67(3):503–9, b) Peng Lin, Jack Ho Wong and Tzi Bun NG; A defensin with highly potent antipathogenic activities from the seeds of purple pole bean, Biosci. Rep. (2010) /30 / 101–109,) Andrew K Joe et.al. Resveratrol Induces Growth Inhibition, S-phase Arrest, Apoptosis, and Changes in Biomarker Expression in Several Human Cancer Cell Lines, Clinical Cancer Research 8, 2002, 893–903, c) Ahmad et.al; Anti-cancer activity of Quercetin, Gallic acid and Ellagic acid against HEPG2 and HCT116 cell lines in vitro. Int J Pharm Bio Sci 2016 Oct ; 7(4): (B) 584 – 592, d) Stephen Barnes, Anticancer Effects of Genistein, Effect of Genistein on In Vitro and In Vivo Models of Cancer, J. Nutr. 1995 125: 3 Suppl 777S–783S, d) Goel A, Aggarwal BB, Curcumin, the golden spice from Indian saffron, is a chemosensitizer and radiosensitizer for tumors and chemoprotector and radioprotector for normal organs. Nutr Cancer. 2010;62(7):919–30. doi:10.1080/01635581.2010.509835.

30. a) Poon, Ivan & Baxter, Amy & Lay, Fung & Mills, Grant & Adda, Christopher & Payne, Jennifer & Phan, Thanh Kha & Ryan, Gemma & White, Julie & Veneer, Prem & van der Weerden, Nicole & Anderson, Marilyn & Kvansakul, Marc & Hulett, Mark. (2014). Phosphoinositide-mediated oligomerization of a defensin induces cell lysis. eLife. 3. e01808, b) Amaral VSGD, Santos SACS, de Andrade PC, et al. Pisum sativum Defensin 1 Eradicates Mouse Metastatic Lung Nodules from B16F10 Melanoma Cells. Int J Mol Sci. 2020;21(8):2662.

31. For IC-50 of standard chemotherapeutics: Kline et.al; Agents to induce apoptosis of p53 mutant, triple-negative human breast cancer cells via activating p73. Breast Cancer Research 2011, 13, b) Young-EE Kwon et.al; In Vitro Histoculture Drug Response Assay and In Vivo Blood Chemistry of a Novel Pt (IV) Compound, K104, Anticancer research 27: 321–326 (2007), c) Fumiyuki Yamasaki, Dongwei Zhang, Chandra Bartholomeusz, Tamotsu Sudo, Gabriel N. Hortobagyi, Sensitivity of breast cancer cells to erlotinib depends on cyclin-dependent kinase 2 activity, Mol Cancer Ther. 2007;6:2168–2177 Fig 1, page, 2171, d) Moon DO, Kim MO, Heo MS, Lee JD, Choi YH, Kim GY. Gefitinib induces apoptosis and decreases telomerase activity in MDA-MB-231 human breast cancer cells. Arch Pharm Res. 2009 Oct 32(10):1351–60, (ranges from 13-46.5 mM for paclitaxel, Cisplatin, Erlotinib, Gefitinib in MDA-MB-231).

32. Tani, K., Murphy, W. J., Chertov, O., Salcedo, R., Koh, C. Y., Utsunomiya, I., Funakoshi, S., Asai, O., Herrmann, S. H., Wang, J. M., Kwak, L. W., Oppenheim, J. J. Defensins Act as Potent Adjuvants That Promote Cellular and Humoral Immune Responses in Mice to a Lymphoma Idiotype and Carrier Antigens. International Immunology 2000. https://doi.org/10.1093/intimm/12.5.691.

33. Gould A, Ji Y, Aboye TL, Camarero JA. Cyclotides, a novel ultra stable polypeptide scaffold for drug discovery. Curr Pharm Des. 2011;17(38):4294–4307.

34. Conibear AC, Rosengren KJ, Daly NL, Henriques ST, Craik DJ. The cyclic cystine ladder in θ-defensins is important for structure and stability, but not antibacterial activity. J Biol Chem. 2013;288(15):10830–10840. doi:10.1074/jbc.M113.451047.

35. Pandurangi, et.al; Drug Development Reports for IND and NDA of AccuTect and NeoTect for FDA and EMEA approval, Schering AG, 2001-2004. b) Taillefer R, Edell S, Innes G, Lister-James J, and the Multicenter Trial Investigators. Acute thromboscintigraphy with 99mTc-apcitide: results of the phase 3 multicenter clinical trial comparing 99mTc-apcitide scintigraphy with contrast venography for imaging acute DVT. J Nucl Med. 2000;41,1214–1223.

36. (1) Abdullah, L. N., Chow, E. K. Mechanisms of Chemoresistance in Cancer Stem Cells. Clinical and Translational Medicine 2013. https://doi.org/10.1186/2001-1326-2-3.

37. For Ceramide See: a) Segui B, Andrieu-Abadie N, Jaffrezou JP, Benoist H, Levade T. Sphingolipids as modulators of cancer cell death: potential therapeutic targets. Biochim Biophys Acta 2006; 1758:2104–20. (b), ASK1 Pathway: Saitoh, M., Nishitoh, H., Fujii, M., Takeda, K., Tobiume, K., Sawada, Y., Kawabata, M., Miyazono, K. & Ichijo, H. Mammalian thioredoxin is a direct inhibitor of apoptosis signal-regulating kinase (ASK)1, (1998) EMBO J. 17, 2596–2606.

38. Simons, A. L., Parsons, A. D., Foster, K. A., Orcutt, K. P., Fath, M. A., Spitz, D. R. Inhibition of Glutathione and Thioredoxin Metabolism Enhances Sensitivity to Perifosine in Head and Neck Cancer Cells. Journal of Oncology 2009. https://doi.org/10.1155/2009/519563.

39. a) Schroeder, B. O., Wu, Z., Nuding, S., Groscurth, S., Marcinowski, M., Beisner, J., Buchner, J., Schaller, M., Stange, E. F., Wehkamp, J. Reduction of Disulphide Bonds Unmasks Potent Antimicrobial Activity of Human β 2-Defensin 1. Nature 2011. https://doi.org/10.1038/nature09674, b) Raghu Pandurangi, Paulmurugan et.al: Restoration of Human Beta Defensin (hBD-1) levels in vivo as immunomodulate for cancer therapy, NIH Report 2015, Immunity, 2020 (submitted).

40. Sugahara, K. N., Teesalu, T., Prakash Karmali, P., Ramana Kotamraju, V., Agemy, L., Greenwald, D. R., Ruoslahti, E. Coadministration of a Tumor-Penetrating Peptide Enhances the Efficacy of Cancer Drugs. Science 2010. https://doi.org/10.1126/science.1183057.

41. Chou TC. Theoretical basis, experimental design, and computerized simulation of synergism and antagonism in drug combination studies. Pharmacol Rev. 2006; 58:621–81.

42. a) Wang, X., Han, W., Yu, Y., Zhu, J., Zhang, R. Doxorubicin-Induced Cardiomyopathy. In Doxorubicin: Biosynthesis, Clinical Uses and Health Implications; 2014. https://doi.org/10.1056/nejm199809243391307, b) Chatterjee, K., Zhang, J., Honbo, N., Karliner, J. S. Doxorubicin Cardiomyopathy. Cardiology. 2010. https://doi.org/10.1159/000265166, c) Renu, K.V.G.A., Tirupathi, T. P., Arunachalam, S. Molecular Mechanism of Doxorubicin-Induced Cardiomyopathy – An Update. European Journal of Pharmacology. 2018. https://doi.org/10.1016/j.ejphar.2017.10.043.

43. (1) Zhou, H. Z., Swanson, R. A., Simonis, U., Ma, X., Cecchini, G., Gray, M. O. Poly(ADP-Ribose) Polymerase-1 Hyperactivation and Impairment of Mitochondrial Respiratory Chain Complex I Function in Reperfused Mouse Hearts. American Journal of Physiology - Heart and Circulatory Physiology 2006. https://doi.org/10.1152/ajpheart.00823.2005.

44. a) Tomas Ganz Defensins: antimicrobial peptides of innate immunity, Nature Reviews Immunology 3, 2003, 710–720, b) Lacerda AF, Vasconcelos ÉAR, Pelegrini PB, Grossi de Sa MF. Antifungal defensins and their role in plant defense. Frontiers in Microbiology. 2014; 5:116.

45. (1) Squires, M. H., Volkan Adsay, N., Schuster, D. M., Russell, M. C., Cardona, K., Delman, K. A., Winer, J. H., Altinel, D., Sarmiento, J. M., El-Rayes, B., Hawk, N., Staley, C. A., Maithel, S. K., Kooby, D. A. Octreoscan Versus FDG-PET for Neuroendocrine Tumor Staging: A Biological Approach. Annals of Surgical Oncology 2015. https://doi.org/10.1245/s10434-015-4471-x.

46. Nahta, R., Esteva, F. J. Herceptin: Mechanisms of Action and Resistance. Cancer Letters. 2006. https://doi.org/10.1016/j.canlet.2005.01.041.

